# NMR metabolomics of symbioses between bacterial vaginosis associated bacteria

**DOI:** 10.1101/2021.11.17.468714

**Authors:** Victoria Horrocks, Charlotte K. Hind, Matthew E. Wand, Joel Chan, Jade C. Hopkins, Georgina L. Houston, Rachel M. Tribe, J. Mark Sutton, A. James Mason

## Abstract

Bacterial vaginosis (BV) is a dysbiosis of the vaginal microbiome, characterised by low levels of lactobacilli and overgrowth of a diverse group of bacteria, and associated with higher risk of a variety of infections, surgical complications, cancer and spontaneous preterm birth (PTB). Despite the lack of a consistently applicable aetiology, *Prevotella* spp. are often associated with both BV and PTB and *P. bivia* has known symbiotic relationships with both *Peptostreptococcus anaerobius* and *Gardnerella vaginalis*. Higher risk of PTB can also be predicted by a composite of metabolites linked to bacterial metabolism but their specific bacterial source remains poorly understood. Here we characterise diversity of metabolic strategies among BV associated bacteria and lactobacilli and the symbiotic metabolic relationships between *P. bivia* and its partners and show how these influence the availability of metabolites associated with BV/PTB and/or pro- or anti-inflammatory immune responses. We confirm a commensal relationship between *Pe. anaerobius* and *P. bivia*, refining its mechanism; *P. bivia* supplies tyrosine, phenylalanine, methionine, uracil and proline, the last of which leads to a substantial increase in overall acetate production. In contrast, our data indicate the relationship between *P. bivia* and *G. vaginalis* strains, with sequence variant G2, is mutualistic with outcome dependent on the metabolic strategy of the *G. vaginalis* strain. Seven *G. vaginalis* strains could be separated according to whether they performed mixed acid fermentation (MAF) or bifid shunt (BS). In co-culture, *P. bivia* supplies all *G. vaginalis* strains with uracil and received substantial amounts of asparagine in return. Acetate production, which is lower in BS strains, then matched that of MAF strains while production of aspartate increased for the latter. Taken together, our data show how knowledge of inter- and intra-species metabolic diversity and the effects of symbiosis may refine our understanding of the mechanism and approach to risk prediction in BV and/or PTB.

## Introduction

Bacterial vaginosis (BV) is regarded as a disruption of the lower genital tract microbiota with a shift from lactobacilli dominance to include a greater proportion of a range of species including members of the genera *Gardnerella, Prevotella, Atopobium, Mobiluncus*, and *Peptostreptococcus* as well as *Sneathia, Leptotrichia, Mycoplasma*, and BV-associated bacterium 1 (BVAB1) to BVAB3.^1^ Despite the lack of consistent aetiology documented in women with BV, vaginal dysbiosis involving a plethora of species, irrespective of whether symptoms of BV are present, promotes local inflammation and is associated with a wide array of health problems.^1^

A specific complication that may be related to BV is a 2-fold increased risk of spontaneous preterm birth (PTB).^2,3^ However, screening for asymptomatic BV in pregnancy in low-risk groups has not aided preterm birth prediction and evidence is insufficient or conflicting even in studies of higher risk groups. Nevertheless, numerous studies have pursued the association between the vaginal microbiome and PTB risk,^5-16^, including our own,^15^ and many of the species identified as associated with higher risk of PTB overlap with those associated with BV.

Changes in microbiota composition are reflected in variations in bacterial derived metabolite profiles,^11,15,17^ which may have functional impact.^18-21^ Consistent with the microbiome studies, elevated vaginal lactate, which is the major product of the lactobacilli, and succinate have been found to be associated with term delivery,^11^ while elevated acetate was subsequently found to be higher in women who delivered preterm compared with term.^17^ A role for these metabolites in BV has also been considered,^18,21^ with two studies agreeing that low lactate and high acetate and propionate are characteristic of BV.^22,23^ Recently, we have shown that combining microbiome and metabolome into composite models has predictive value for preterm birth.^15^ A composite of metabolites which include lactate and acetate but also, aspartate, leucine, tyrosine and betaine associated with risk of PTB < 37 weeks while risk of PTB < 34 weeks was identified by a composite comprising *L. crispatus, L. acidophilus*, glucose and, again, aspartate.

Although multiple studies have identified *Prevotella* spp. as being associated with both BV, and preterm birth,^9,12,13,15^ their presence has not been found to be predictive of PTB.^15^ However, their residence within the vagina correlates with that of a number of other bacteria including *Gardnerella vaginalis*,^15,16^ and *P. bivia* is known to enjoy symbiotic interactions with both *Peptostreptococcus anaerobius* and *G. vaginalis*.^24-26^ Two groups have found an association between preterm birth and *G. vaginalis*,^7,9,16^ but its presence alone does not predict PTB. There is though, reason to consider whether the substantial diversity of *G. vaginalis* affects the ability to establish its functional role(s) in both BV and preterm birth.^27^ Studies of microbial communities often sequence and quantify specific marker genes and cluster such sequences into Operational Taxonomic Units (OTUs). Although such OTUs have been generally shown to have high levels of ecological consistency,^28^ and the approach remains popular and useful, there remains the possibility that functionally relevant differences in bacterial behaviour are obscured by this approach. Indeed, in one study that confirmed an association between *G. vaginalis* and preterm birth, high-resolution statistical bioinformatics was used to detect nine unique *G. vaginalis* 16S rRNA sequence variants and this revealed that only one of three *G. vaginalis* clades was responsible for the association of the genus with PTB.^9^ Strain level profiling has also helped improve understanding of species co-occurrence profiles.^16^

In addition, the role of the otherwise dominant lactobacilli may also be critical in defining PTB risk, with *Lactobacillus crispatus* dominance frequently associated with term delivery.^9,10,13,15,16^ The picture for *L. iners* is less clear. One study showed an association with PTB,^10^ but two subsequent studies found none.^9,15^ Instead they found frequent co-existence of *L. iners* with *G. vaginalis*,^9^ which contrasts with *L. crispatus* where an exclusionary relationship with *G. vaginalis* is found,^9,16^ or positive correlation with BV associated bacteria including *P. bivia*.^15^

Given the valuable utility of the NMR metabolomics approach for identifying risks associated with vaginal dysbiosis and predicting PTB, and associations with differing microbiome states likely to have functional impact, there is an unmet need to understand bacterial contributions to the vaginal metabolome in more detail. To this end, we aimed to establish a mechanistic basis for a mutualistic symbiotic relationship between *P. bivia* and *G. vaginalis* and contrast this with the commensal relationship between *P. bivia* and *Pe. anaerobius*. We characterise the diverse metabolic strategies of a panel of *G. vaginalis* isolates and determine how this influences symbiosis with *P. bivia*. In addition, we compare metabolism across a panel of lactobacilli to highlight that variation in metabolic strategy is not limited to BV/sPTB associated bacteria and that the metabolite background will likely vary according to microbiome community state type (CST).^5^ The information provided by the present study suggests ways of refining prediction models that include metabolite data and gives insight into how bacterial metabolism and symbiosis influence each other, with implications for functional impact and clinical outcomes.

### Experimental procedures

#### Isolates

*Gardnerella vaginalis* 11292, 10915 and 10287, *Peptostreptococcus anaerobius* 11460, *Prevotella bivia* 11156, *Atopobium vaginae* 13935 and *Mobiluncus curtisii* 11656 were obtained from the National Collection of Type Cultures (NCTC). All other bacteria were isolated from swabs collected from pregnant women recruited with informed written consent via the INSIGHT study (NHS Human Research Authority, London - City and East Research Ethics Committee 13LO/1393) or from Salisbury District Hospital (SDH) microbiology lab. Samples from SDH were received from the microbiology laboratory following diagnostic testing. All identifiers were re-moved by the diagnostic laboratory’. The swabs were maintained at ambient temperature during transport in liquid amies buffer and were used immediately or frozen at −80°C until use. 100 µl of the buffer solution was either plated onto tryptic soy agar (TSA) and De Man, Rogosa and Sharpe (MRS) agar plates and incubated at 37°C for 48 hours under aerobic condition or plated onto TSA, MRS agar and Columbia blood agar (CBA), containing 5% defibrinated sheep’s blood (Oxoid), and incubated at 37°C for 48 hours under anaerobic conditions as outlined below. Single colonies were streaked to purity and identified using MALDI-TOF spectrometry (MALDI Biotyper®, Bruker Daltonics GmbH & Co. KG, DE).

#### Bacterial culturen

All *G. vaginalis* isolates, *Pe. anaerobius, P. bivia, M. curtisii* and *A. vaginae* were plated onto CBA (Oxoid, Hampshire, UK) containing 5% defibrinated sheep’s blood (Oxoid) and incubated at 37°C for 48 hours under anaerobic conditions generated using Thermo Scientific™ Oxoid™ AnaeroGen™. *L. iners* was plated under the same conditions for 72 hours. All other *Lactobacillus* species were plated onto MRS agar (Sigma Aldrich) and incubated at 37°C for 48 hours under anaerobic conditions. For initial overnight cultures a 1 µl loop of culture was used to inoculate 5 ml of brain-heart infusion (BHI) media with 5% horse serum and incubated at 37°C for 48 hours under anaerobic conditions without shaking. For monoculture samples, 50 µl of overnight culture was added to 5 ml of fresh BHI with 5% horse serum and incubated at 37°C for 48 hours under anaerobic conditions without shaking. For coculture of *P. bivia* with *G. vaginalis* or *Pe. anaerobius*, from overnight cultures, a 1:1 mix of each species was used to inoculate 5 ml of fresh BHI with 5% horse serum and incubated at 37°C for 48 hours under anaerobic conditions without shaking.

#### MIC testing

The minimum inhibitory concentrations (MICs) were measured using a broth microdilution method in polypropylene plates (Greiner). From an overnight culture in BHI 100 µl of bacterial culture totalling an OD_600_ of 0.1 was added to 100 µl of BHI media containing antibiotic. After 48 hours of incubation at 37°C under anaerobic conditions the optical density at a wavelength of 600 nm was read. The lowest concentration of antibiotic where there was no growth (OD_600_ < 0.1) determined the MIC.

#### NMR metabolomics

For preparation of samples to be used in metabolomics bacterial cultures were pelleted by centrifuge at 5000 rpm at 4°C. Supernatant was filtered with 0.22 µm membrane to remove any bacterial cells and large debris and were stored at −80°C until use. To aid suppression of the water signal and deuterium lock and act as an internal reference, 60 µl of D_2_O + 3-(trimethylsilyl)propionic-2,2,3,3-d4 acid sodium salt (TSP-d4) was added to 570 µl of filtered supernatant. The pH of all samples was adjusted using NaOH to within 0.2 pH units of the BHI media control. ^1^H NMR spectra were recorded on Bruker 600 MHz Bruker Avance III NMR spectrometer (Bruker BioSpin, Coventry, United Kingdom) equipped with a 5 mm ^1^H, ^13^C, ^15^N TCI Prodigy Probe and a cooled sample changer with all samples kept at 4 °C. The 1D spectra were acquired under automation at a temperature of 298 K using Carr-Purcell-Meiboom-Gill presaturation (CMPG) pulse sequence (cpmgrp1). The parameters of spectra acquisition are 32 transients, a spectral width of 20.83 ppm and 65,536 datapoints. For assignment of metabolite peaks additional spectra, Total correlation spectroscopy (TOCSY), ^1^H-^13^C heteronuclear single quantum correlation spectroscopy and J-resolved spectroscopy (JRES), were acquired from a pooled sample containing a small volume of all samples. Resonance positions are quoted in ppm with respect to the methyl peak of TSP-d4 at 0.0 ppm.

All spectra were Fourier transformed in Bruker software and adjusted using automatic baseline correction and phasing in Bruker TopSpin 4.1.3. Multiple databases were used for the assignment of metabolites; Chenomx NMR suite software (Chenomx Inc, Canada), Human Metabolome Database (HMDB) and Biological Magnetic Resonance Data Bank (BMRB).^29^ To convert NMR intensity to mM concentration the Chenomx software programme was used calibrated to the concentration of TSP-d4 present in the sample. For multivariate analysis the intensity of all samples was normalised using probabilistic quotient normalisation (PQN).^30^ For visualisation of data, python packages numpy, matplotlib, seaborn, pandas and scipy were used.

#### Sequencing

All isolates identified as *G. vaginalis* from MALDI-TOF were also confirmed through whole genome sequencing. DNA was extracted from overnight culture in BHI using the GenElute™ Bacterial Genomic DNA Kits (Sigma Aldrich). DNA was tagged and multiplexed with the Nextera XT DNA kit (Illumina, San Diego, US) and sequenced by Public Health England Genomic Services and Development Unit, (PHE-GSDU) on an Illumina (HiSeq 2500) with paired-end read lengths of 150 bp. A minimum 150 Mb of Q30 quality data were obtained for each isolate. FastQ files were quality trimmed using Trimmomatic^31^. SPAdes 3.1.1 was used to produce draft chromosomal assemblies, and contigs of less than 1 kb were filtered out^32^. Whole genome alignment and phylogenetic tree generation were performed using progressive alignment in Mauve Version 20150226 build 10. Tree visualisation was performed in FigTree Version 1.4.3.

## Results

To better understand the contribution of different bacteria to the vaginal metabolome in eubiosis and dysbiosis a panel of lactobacilli and BV associated isolates was assembled. Whole genome sequencing of seven *G. vaginalis* strains included reference strains from the NCTC and new isolates from vaginal swabs, enables them to be assigned to Clades^9,16,33^ or subgroup^34^ and identifies genes for sialidase and vaginolysin (Table 1). Though expression was not tested, all isolates carry the genes coding for sialidase and vaginolysin. Strains KC1 and KC2 have Type 1B vaginolysin while the remainder have Type 1A.^35^ Six of the seven strains are members of Clade 1/Subgroup C/Clade GV2a, corresponding to sequence variant G2 strains which have been shown to drive observed associations with PTB.^9^ The remaining isolate, KC1, is a member of Clade 3/Subgroup D/Clade GV1b. Tested for susceptibility to the main antibiotics used for BV, two isolates, KC1 and KC3, are found to be resistant to metronidazole and tinidazole. All isolates are sensitive to clindamycin and erythromycin.

**Table 1.**
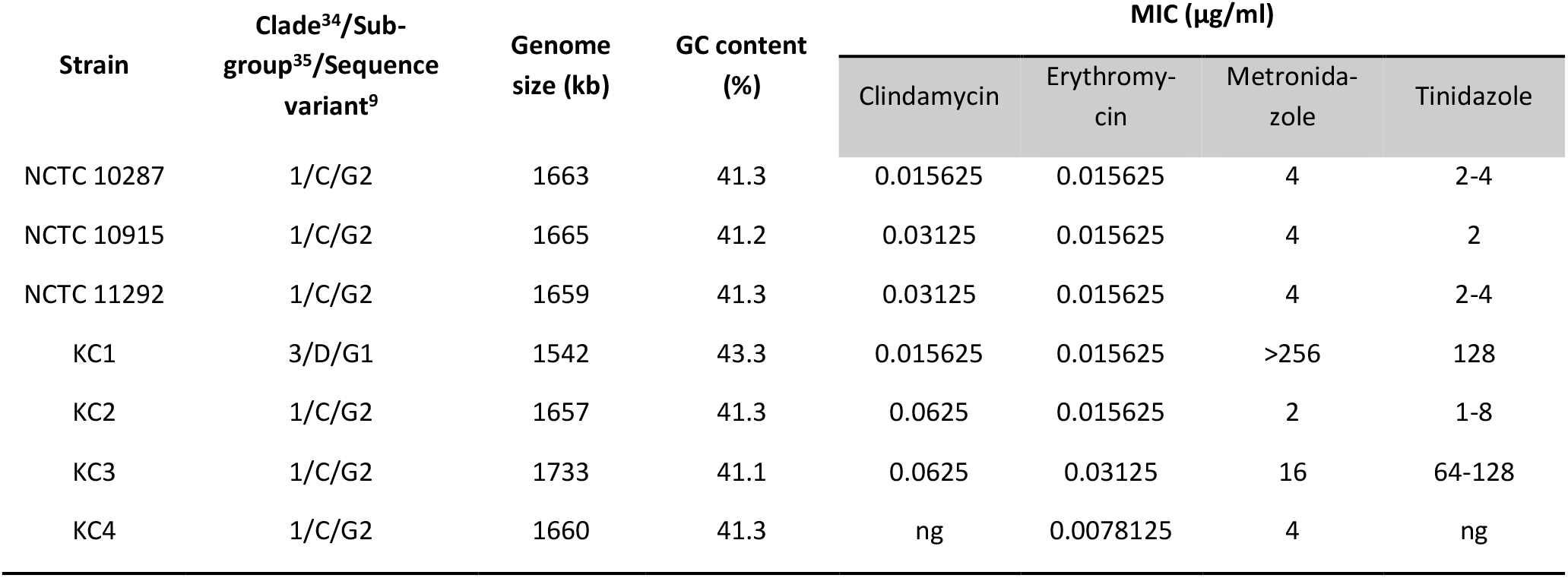
*G. vaginalis* strain characteristics. All strains are positive for the genes encoding sialidase and vaginolysin. Concordant MICs are reported from three independently repeated experiments. ng = no growth.

| Strain | Clade<sup>34</sup>/Sub-group<sup>35</sup>/Sequence variant<sup>9</sup> | Genome size (kb) | GC content (%) | MIC (µg/ml) |  |  |  |
| --- | --- | --- | --- | --- | --- | --- | --- |
|  |  |  |  | Clindamycin | Erythromycin | Metronidazole | Tinidazole |
| NCTC 10287 | 1/C/G2 | 1663 | 41.3 | 0.015625 | 0.015625 | 4 | 2-4 |
| NCTC 10915 | 1/C/G2 | 1665 | 41.2 | 0.03125 | 0.015625 | 4 | 2 |
| NCTC 11292 | 1/C/G2 | 1659 | 41.3 | 0.03125 | 0.015625 | 4 | 2-4 |
| KC1 | 3/D/G1 | 1542 | 43.3 | 0.015625 | 0.015625 | >256 | 128 |
| KC2 | 1/C/G2 | 1657 | 41.3 | 0.0625 | 0.015625 | 2 | 1-8 |
| KC3 | 1/C/G2 | 1733 | 41.1 | 0.0625 | 0.03125 | 16 | 64-128 |
| KC4 | 1/C/G2 | 1660 | 41.3 | ng | 0.0078125 | 4 | ng |

### Overview of bacterial metabolism in BHI and identification of major metabolic strategies for BV associated bacteria

Analysis of BHI spent culture allows comparison of the overall metabolic strategy for each of the BV associated bacteria but also comparison (Fig. 1) of the relative amounts of key metabolites that define the vaginal chemical environment and/or have been associated with BV and/or PTB (Fig. 2; S1). The NMR metabolomic approach clearly identifies the pyruvate and/or glucose fermentative strategies of *A. vaginae, Pe. anaerobius* and the seven *G. vaginalis* isolates. The seven *G. vaginalis* isolates can be distinguished from each other and classified according to whether they use the bifid shunt (BS) alone, producing lactate and acetate from glucose,^36^ or mixed acid fermentation (MAF) producing lactate and acetate but also formate and ethanol and consuming pyruvate in addition to glucose (Fig. 1A; 2A/B/E/F). *G. vaginalis* 10287 and 10915 are hence classified as using BS alone while the remainder all use MAF.

**Figure 1.**
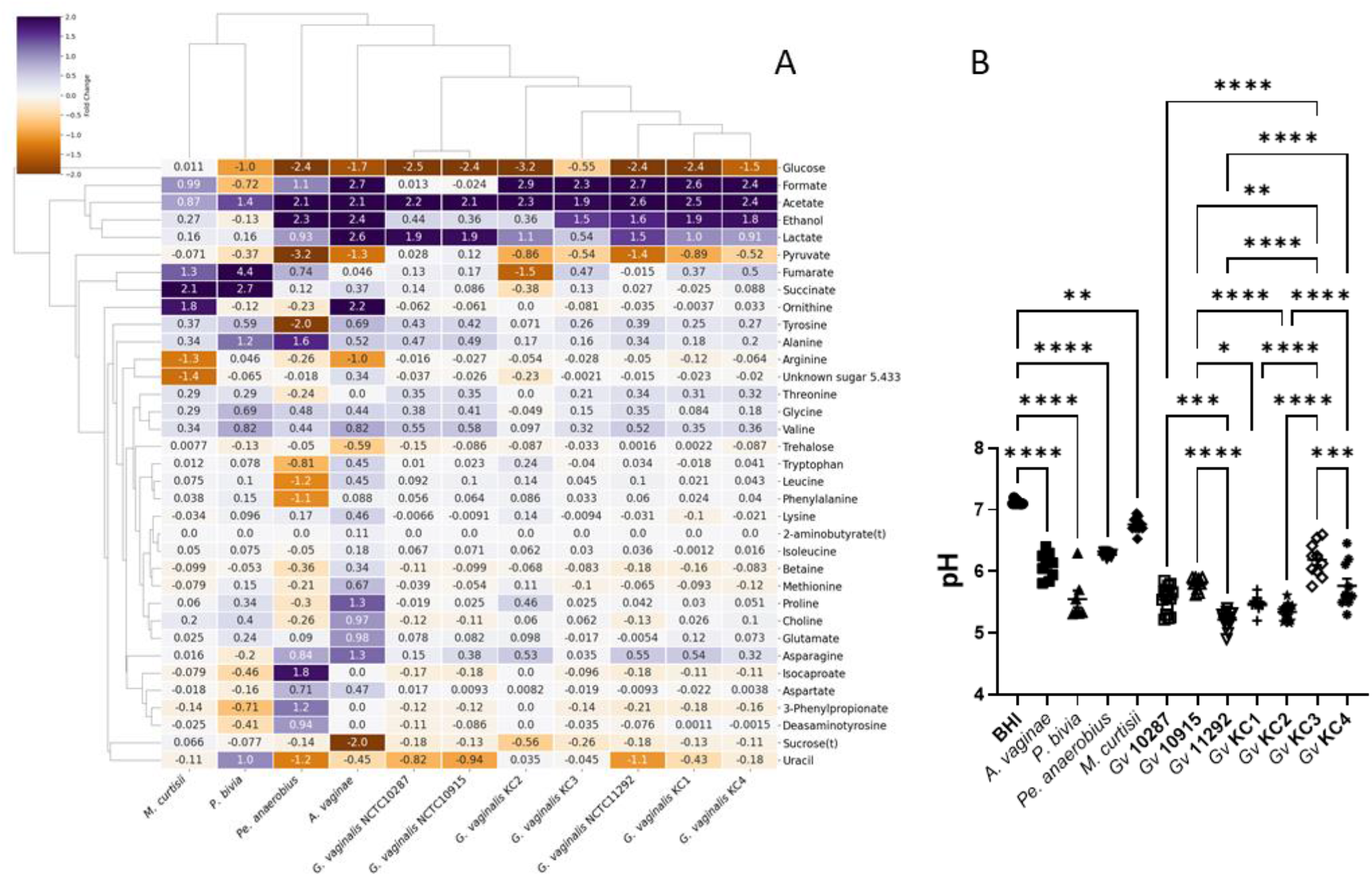
Diversity of bacterial vaginosis associated bacteria metabolism, when cultured in brain heart infusion. The heatmap (A) compares the metabolite fold change from ^1^H NMR of spent culture media and enables the major metabolites produced by each isolate and the key differences in metabolic strategy to be revealed. The resulting acidification of the spent culture media accordingly varies (B). Comparisons are shown between fresh BHI and the five non-*G. vaginalis* conditions and between each of the *G. vaginalis* strains, as determined by One-way ANOVA with Tukey correction for multiple comparisons. * *p* < 0.05, ** *p* < 0.01, *** *p* < 0.001, **** *p* < 0.0001. All *G. vaginalis* strains acidify the media (*p* < 0.0001).

**Figure 2.**
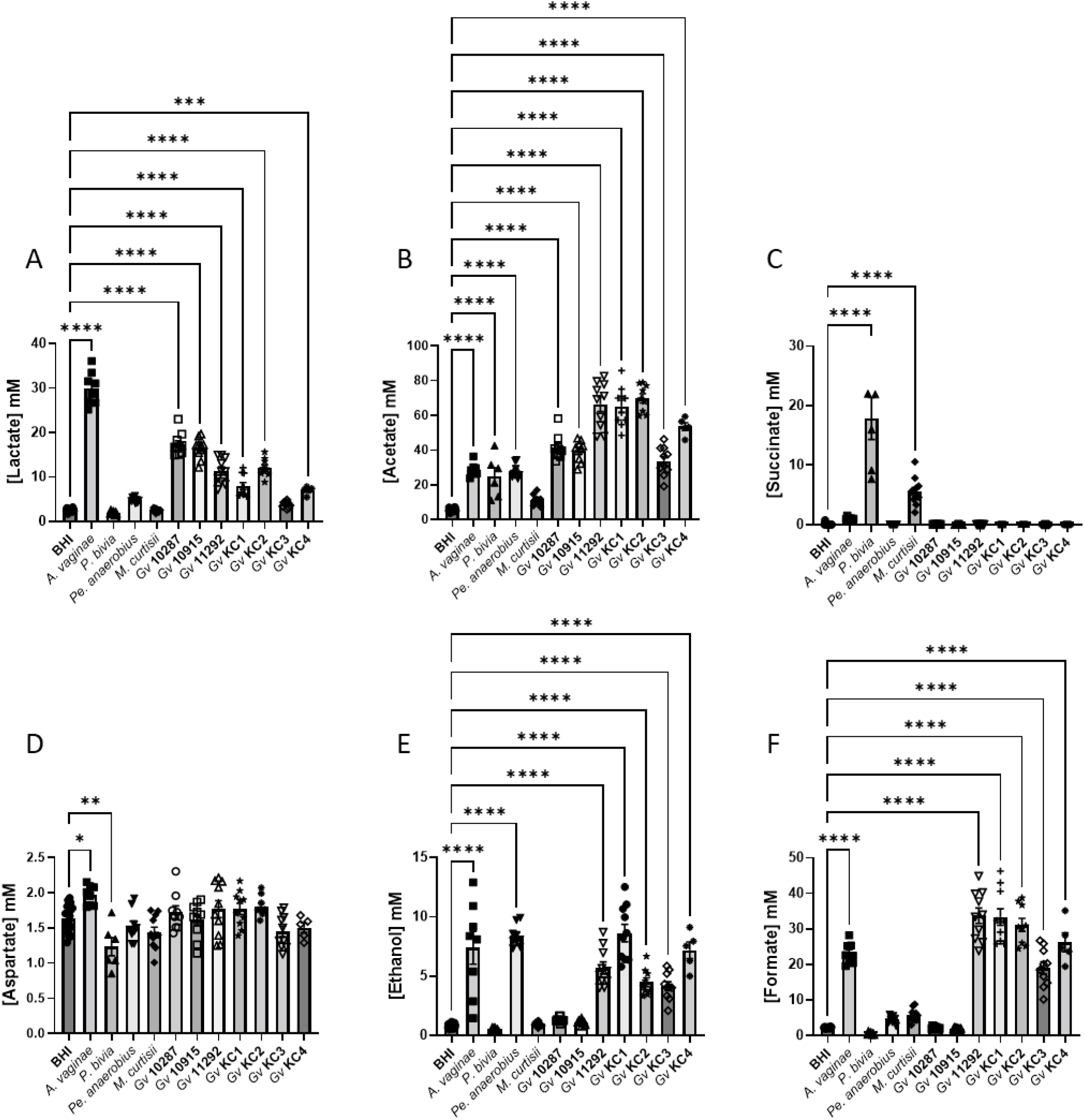
Production of organic acids, aspartate and ethanol by bacterial vaginosis associated bacteria in BHI. Comparisons are made between BHI and each spent culture as determined by One-way ANOVA with Tukey correction for multiple comparisons for the main products of fermentation and/or those involved in anaerobic respiration. Comparisons for other metabolites presented in Fig. S1. Only pairwise comparisons where *p* < 0.05 are shown. * *p* < 0.05; ** *p* < 0.01; *** *p* < 0.001; **** *p* < 0.0001.

*M. curtisii* is known to be capable of using trimethylamine oxide (TMAO) as an electron donor for anaerobic respiration, producing trimethylamine (TMA).^37^ In BHI it also conducts anaerobic respiration, but the production of succinate (Fig. 1; 2C) is suggestive of fumarate acting as the electron donor in place of TMAO which is absent. *M. curtisii* is known also to consume arginine to produce ornithine, citrulline and ammonia,^38^ and both it and *A. vaginae* do this also in BHI (Fig. 1A; S1E/T). *P. bivia* characteristically also produces succinate via anaerobic respiration but also ferments glucose to acetate,^39^ and this is observed in BHI alongside avid consumption of asparagine (Fig. 1A; 2B/C; S1D). *P. bivia* notably excretes a variety of metabolites that are not produced at the same levels or at all, and are often consumed, by the other BV associated bacteria. These include succinate and fumarate and alanine, glutamate, glycine, methionine, phenylalanine, proline, valine and uracil (Fig. 1A; S1C/H/J/L/M/N/O/P/R/S).

The result of these differing metabolic strategies is, in every case, an acidification of the spent BHI culture but this is relatively modest for *M. curtisii, Pe. anaerobius* and *A. vaginae* compared with that observed for the seven *G. vaginalis* strains and *P. bivia* (Fig. 1B).

Considering the lactobacilli, four species are considered obligate homofermentative (*L. acidophilus, L. crispatus, L. gasseri* and *L. iners*) using the Embden-Meyerhof-Parnas (EMP) pathway to make lactate (both D-lactate and L-lactate with the exception of *L. iners* that makes only L-lactate), two species are considered facultative heterofermentative making lactate (L-lactate for *L. rhamnosus* and D-lactate for *L. jensenii*) and acetate and one species, *L. fermentum*, is obligate heterofermentative producing lactate, acetate and ethanol as well as CO_2_.^40^ The present NMR results are consistent with this with all lactobacilli producing lactate (Fig. S2A; S3A), only *L. fermentum* producing substantial quantities of ethanol (Fig. S2A; S3E) and only *L. rhamnosus* producing substantial amounts of formate (Fig. S2A; S3F). Consistent with genome sequence studies, which showed a lack of enzymes to produce acetate,^41,42^ *L. iners*, is the only *Lactobacillus* in this study that does not produce any acetate in BHI; acetate production by the other lactobacilli varies considerably (Fig. S2A; S3B). The lactobacilli can be further distinguished, notably at strain level for *L. crispatus*, by differing consumption of pyruvate, asparagine, arginine, glycine, lysine and proline (Fig. S4B/D/J/K/M/P/R) and production of alanine, valine, isoleucine and uracil (Fig. S4K/S/T/U). Acidification of the spent culture media is likely limited by the relatively low glucose concentration in BHI but the greatest acidification is achieved by *L. acidophilus* (significantly more than all except *L. crispatus* 2), which also produces more lactate than any of the other strains (*p* < 0.05) (Fig. S2B; S3A).

We have shown previously that lower lactate and higher acetate were associated with increased risk of PTB < 37 weeks (odds ratios respectively 0.432 and 1.610).^15^ As expected, and again despite the relatively low concentration of glucose in BHI, the lactobacilli produce a final lactate concentration of between 30 and 45 mM in spent BHI culture (Fig. S3A), which substantially exceed production by BV associated bacteria (Fig. 2A). Of note however is that *A. vaginae* spent culture is enriched with around 27 mM lactate and the two BS *G. vaginalis* produce substantially more lactate than the five MAF *G. vaginalis* strains (*p* < 0.0001) (Fig. 2A). Except for *L. iners*, acetate is produced by lactobacilli in BHI to achieve final concentrations ranging from 5 mM to 21 mM (Fig. S3). Similar levels of acetate production are achieved by *A. vaginae, P. bivia* and *Pe. anaerobius* but this is dwarfed by production by *G. vaginalis* with BS strains attaining c 35 mM and MAF strains as much as 65 mM. Although succinate secretion is a hallmark of anaerobic respiration and concentrations of nearly 18 mM are achieved in *P. bivia* spent culture (Fig. 2C), small amounts of this dicarboxylate (1-4 mM) are also detected in all lactobacilli spent cultures with the ex-ception again of *L. iners* (Fig. S3C).

Higher aspartate has previously been associated with increased risk of PTB < 37 and < 34 weeks (odds ratios respectively 1.675 and 1.768).^15^ Seven of the nine lactobacilli strains produce this, but this is very modest with spent culture enriched by a maximum of 1.2 mM aspartate (Fig. S3D). In monoculture, none of the *G. vaginalis* strains produce aspartate but modest amounts are produced by *A. vaginae* and it is consumed by *P. bivia* (Fig. 2D). We have reported higher glucose associated with increased risk of PTB < 34 weeks (odds ratio 1.269).^15^ Almost all glucose in BHI is consumed by both lactobacilli and BV associated bacteria, with the exception of *M. curtisii* (Fig. S1A/S4A). *G. vaginalis* KC3 did not consume all glucose in this first study but subsequently it grew well, and its consumption matched that of the other *G. vaginalis* strains (Fig. S1A/S7A). In contrast, pyruvate available in BHI is not universally consumed (Fig. S1B; S4B). *A. vaginae, Pe. anaerobius, L. crispatus* 1 and *L. fermentum* consume all pyruvate available while the remaining lactobacilli and BV associated bacteria, except for *G. vaginalis* 10287 and 10915, consume some but not all. *G. vaginalis* 10287 and 10915 secrete modest amounts of pyruvate into the spent culture (*p* < 0.05).

Higher leucine and betaine and lower tyrosine have also been associated with increased risk of PTB< 37 weeks (odds rations respectively, 3.118, 1.365 and 0.023).^15^ None of the lactobacilli or BV associated bacteria in the present study produce leucine when cultured in BHI though it is avidly consumed by *P. bivia* (Fig. S1G; S4G). Tyrosine is produced in modest amounts by six of the lactobacilli isolates, most notably by *L. acidophilus, L. gasseri* 1 and 2, and most *G. vaginalis* strains as well as *A. vaginae, P. bivia* and *M. curtisii* (Fig S1C). It is consumed avidly by *Pe. anaerobius* (Fig. S1C). With the exception of *Pe. anaerobius, L. crispatus* 1 and *L. iners*, where there is modest consumption, the concentration of betaine does not change in the spent culture of either the lactobacilli or the BV associated bacteria (Fig. S1E; S4H). Similarly, with the exception of *A. vaginae*, changes in choline concentrations are minimal (Fig S1F; S4F).

### Symbiosis between *P. bivia* and *Pe. anaerobius* influences production of key PTB markers

^1^H NMR of the spent culture from *P. bivia, Pe. anaerobius* and a 1:1 co-culture reveals that combining the two species leads to a substantial adjustment in the levels of metabolites that have previously been associated with PTB and/or shown utility in predicting patient outcomes. In the spent BHI media, even though relative abundance could not be enumerated by plating, there is clear evidence from production and consumption of species-specific metabolites that both species proliferate (Fig. 3). In monoculture, only *P. bivia* consumes asparagine and produces butyrate, fumarate and succinate and this is observed also in co-culture although succinate production is reduced (*p* < 0.0001) (Fig. S5A-D). Similarly, *Pe. anaerobius* is known to have a characteristic organic acid production profile,^43^ and in monoculture, of the two species, only *Pe. anaerobius* consumes phenylalanine, proline, tyrosine, uracil, lysine, methionine, choline and leucine (Fig. S5J-Q) and produces ethanol, 4-methylpentanoate (isocaproate), 3-(4-hydroxyphenyl)propanoate (desaminotyrosine/phloretic acid), 3-phenylpropionate (hydrocinnamate) and 5-aminopentanoate (aminovalerate) (Fig. S5E-I). With the possible exception of choline consumption, this is also observed in coculture, with increased production observed for all five of its specific products.

**Figure 3.**
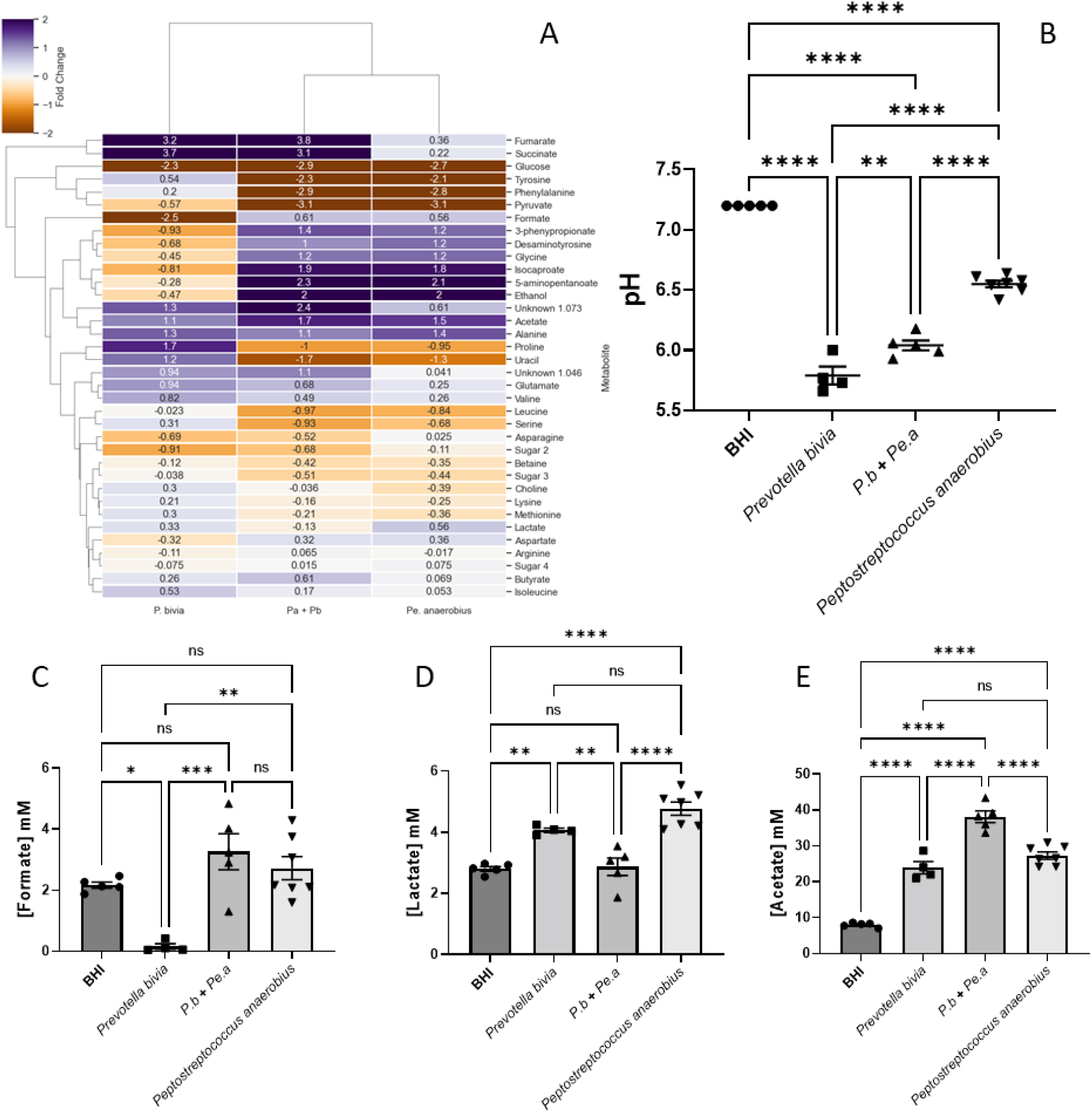
Commensal symbiosis of *P. bivia* and *Pe. anaerobius* in BHI generates a distinct chemical environment. The heatmap compares the metabolite fold change from ^1^H NMR of spent culture media and enables, the major metabolites produced by each isolate and the key differences in metabolic strategy to be revealed (A). The pH of the three spent cultures is compared with fresh BHI (B). Levels of formate (C), lactate (D) and acetate (E) in spent culture are shown relative to fresh BHI. Further metabolites are shown in Fig. S5 and S6. Comparisons are shown between all conditions, as determined by Oneway ANOVA with Tukey correction for multiple comparisons. * *p* < 0.05, ** *p* < 0.01, *** *p* < 0.001, **** *p* < 0.0001.

Previously, a commensal symbiosis between *P. bivia* and *Pe. anaerobius* has been demonstrated and ascribed to use of amino acids, by *Pe. anaerobius*, that were secreted by *P. bivia*.^24^ Here, ^1^H NMR identifies enrichment of BHI media with tyrosine, phenylalanine, proline, methionine, alanine, glutamate, glycine, isoleucine, valine and also choline and uracil (Fig. S5/S6). Of these, *Pe. anaerobius* avidly consumes tyrosine, phenylalanine, proline and uracil, and modestly consumes methionine and possibly choline (Fig. S5). Levels of alanine, glutamate, glycine, isoleucine and valine are also lower in the co-culture spent media than that of *P. bivia* but, since these are available in BHI normally and are not consumed in *Pe. anaerobius* monoculture, it is assumed that this reduction can also be accounted for by a lower overall growth of *P. bivia* in the combination relative to monoculture (Fig. S6).

While the benefits of co-culture to *Pe. anaerobius* appear manifold, the reverse is not true for *P. bivia* and ^1^H NMR does not detect any metabolites produced by *Pe. anaerobius* that are consumed by *P. bivia*. This supports the previous finding of a commensal relationship between the two organisms.^24^ There is one possible caveat to this in that, while no effect of *Pe. anaerobius* conditioned media on *P. bivia* growth was observed previously,^24^ here we find that *P. bivia* metabolism is likely altered by co-culture with *Pe. anaerobius*. First, while production of *Pe. anaerobius* specific metabolites is increased in co-culture relative to monoculture, the same is true for *P. bivia* only for butyrate (*p* = 0.0295), with less succinate, alanine, glutamate, glycine and valine than might be expected. Second, the total consumption of formate by *P. bivia* in monoculture is not observed in co-culture (Fig. 3C) while lactate, produced by both species in monoculture, is no more abundant in the co-culture spent media than in fresh BHI (Fig. 3D) even though acetate production almost doubles (Fig. 3E). Both formate and lactate are potential electron donors for anaerobic respiration and the NMR analysis provides evidence for a switch in electron donor, from formate to lactate, by *P. bivia* when *Pe. anaerobius* is present.

The spent culture media pH will be affected by the production/consumption of a range of organic and amino acids. Although acidification of spent culture will be limited due to the relatively low levels of glucose interactions between these two species will affect the acidity of the environment (Fig. 3B). Despite production of acetate (pKa 4.76) and lactate (pKa 3.86), acidification by *Pe. anaerobius* is relatively modest with a reduction by only 0.65 pH units. In contrast, both the spent *P. bivia* monoculture and co-culture are reduced by over one pH unit (respectively 1.41 and 1.16). In both cases substantial amounts of succinate (pKa 4.2, 5.6) are produced (20 mM vs 9.5 mM for monoculture vs co-culture). More acetate is produced in the co-culture (30.1 vs 15.9 mM) but there is no net lactate production. These effects combine to ensure that the spent co-culture pH is a little higher than that of the *P. bivia* monoculture but substantially lower than that corresponding to *Pe. anaerobius*.

#### Symbiosis between *P. bivia* and *G. vaginalis* is strain and metabolic strategy dependent

Five *G. vaginalis* strains (10287, 10915, 11292, KC1 and KC3), representing both BS and MAF strategies, were selected for co-culture experiments with *P. bivia* NCTC 11156. With the exception of KC1, the only strain in the present study not of sequence variant G2,^9^ positive correlations were detected between the number of CFU identified for either species when plated following co-culture in BHI (Fig. 4B, Table 2), with the strongest positive relationship found for *G. vaginalis* 10287, one of the BS strains.

**Table 2.** *P. bivia* vs *G. vaginalis* co-culture correlation. Relationship between CFU counts for each species as a function of *G. vaginalis* isolate as determined by parametric Pearson or non-parametric Spearman correlation coefficients. Only for KC1 is a negative correlation between the two species found while positive correlations exist for the remaining four isolates.

| *G. vaginalis* strain | Number of XY<br>pairs | Spearman |  | Pearson |  |
| --- | --- | --- | --- | --- | --- |
|  |  | r | P (two-tailed) | r | P (two-tailed) |
| NCTC 10287 | 16 | 0.7917 | 0.0005 | 0.7781 | 0.0004 |
| NCTC 10915 | 9 | 0.6442 | 0.0694 | 0.5419 | 0.1318 |
| NCTC 11292 | 22 | 0.5457 | 0.0086 | 0.5133 | 0.0146 |
| KC1 | 10 | -0.4768 | 0.1645 | -0.6896 | 0.0274 |
| KC3 | 19 | 0.5947 | 0.0072 | 0.6392 | 0.0032 |

**Figure 4.**
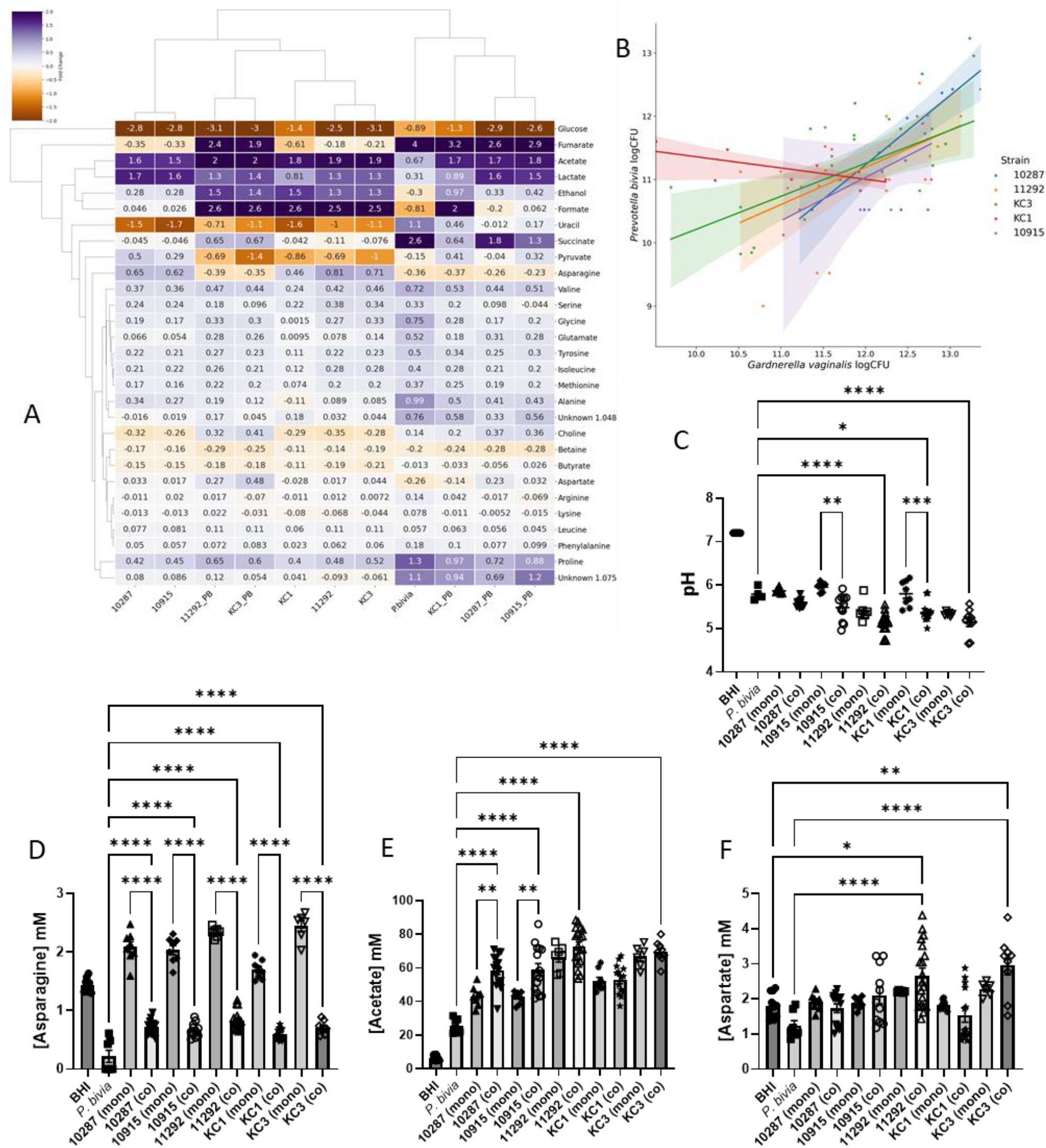
Co-culture of *Prevotella bivia* NCTC 11156 and a panel of *Gardnerella vaginalis* isolates. A heatmap shows the relationship between metabolite fold-changes detected by ^1^H NMR of spent cultures (A). The correlation between CFU for *P. bivia* and each *G. vaginalis* partner is shown for five co-cultures (B); Spearman and Pearson r are provided in Table 1. Spent culture pH for mono- and co-cultures as well as fresh BHI (C). *G. vaginalis* supplies *P. bivia* with asparagine (D). Acetate levels increase when bifid shunt *G. vaginalis* (10287 and 10915) are co-cultured with *P. bivia* (E). Symbiosis between *P. bivia* and MAF *G. vaginalis* strains produces aspartate (F). Comparisons are shown between each co-culture and the corresponding mono-cultures (C-E) and also fresh BHI (F) as determined by One-way ANOVA with Tukey correction for multiple comparisons. Only *p* < 0.05 shown; * *p* < 0.05, ** *p* < 0.01, *** *p* < 0.001, **** *p* < 0.0001. Comparisons for further metabolites shown in Fig. S7.

The Spearman and Pearson r for KC1 are both negative indicating that when *G. vaginalis* KC1 grew well, *P. bivia* did not, and vice versa. This is manifested in the metabolomics analysis where levels of some metabolites, known to be produced by *P. bivia*, notably succinate, fumarate, proline, uracil and alanine are highly variable (Fig. S7D-F, J, K). There is some explanation for this phenomenon in the metabolomics data (Fig.4A). Notably, KC1 may be the only one of the five *G. vaginalis* strains that is not capable of adequately supplying asparagine to *P. bivia* (Fig. 4D). As noted above, *P. bivia* avidly consumes asparagine since this can be used to produce aspartate and, in turn, fumarate which is an important electron acceptor anaerobic respiration. Asparagine is produced in substantial amounts by all *G. vaginalis* isolates (*p* < 0.0001), with the exception of KC1 (*p* = 0.0333). The two BS strains increase the availability of asparagine by 42% (10915) and 45% (10287). This is modest when compared with MAF strains 11292 and KC3 which respectively increase the availability of asparagine by 63% and 70%, such that approximately double the amount of asparagine that is consumed by *P. bivia* in monoculture is available in co-culture. In contrast, KC1 only increases the amount available by 18.5%.

While supply of asparagine from *G. vaginalis* to *P. bivia* is observed for both MAF and BS strains, a further means by which BS strains, but not MAF strains, may supply *P. bivia* is also apparent. Unlike the BS *G. vaginalis* strains, *P. bivia* and all three MAF *G. vaginalis* strains consume pyruvate from BHI (Fig. S7C). With two species growing together the metabolite data for co-culture has greater variance but considering just the data from monocultures (as above) indicates that some pyruvate is likely secreted from 10287 (*p* = 0.0035) and 10915 (*p* = 0.0046). As such the BS strains differ from the MAF strains in that they avoid competition with *P. bivia* for pyruvate and, likely, may supply it in co-culture.

As noted above, in monoculture the MAF strains 11292 and KC3 (*p* < 0.0001) and KC1 (*p* < 0.05) produce more acetate than the BS strains 10287 and 10915 but less lactate. In co-culture however acetate produced by *P. bivia*/10287 and *P. bivia*/10915 increases by 42-45% over that produced by *G. vaginalis* alone while the corresponding figure for the MAF strains is between 2 and 11%. Lactate production is largely unchanged in co-culture for any of the strains. Co-culture with *P. bivia* therefore has the potential to substantially increase overall acetate levels and change the acetate/lactate ratio when BS strains are present but not MAF strains. Further, while *P. bivia* was confirmed to consume formate, ethanol and aspartate by spiking experiments (Fig. S8) there is insufficient evidence here that production of these metabolites by MAF *G. vaginalis* provides substantial benefit for *P. bivia* with no apparent consumption of these metabolites in the respective co-cultures (Fig. S7G, H; Fig. 4F). Indeed, while both 11292 (*p* = 0.015) and KC3 (*p* = 0.008) produce aspartate in monoculture, the amount found in the spent co-culture media is increased respectively 2-and 3-fold (Fig. 4F). As previous work has indicated *P. bivia* supplies ammonia to *G. vaginalis*,^25^ this suggests that MAF *G. vaginalis* might be performing a detoxification role by consuming both ammonia and fumarate (Fig. S7E), secreted by *P. bivia*, to produce aspartate.^44^

While the symbiotic relationship between *P. bivia* and *Pe. anaerobius* is commensal in BHI, we suggest here that the relationship between *P. bivia* and *G. vaginalis* is mutualistic since, as well as the presumed supply of ammonia and fumarate, we show *P. bivia* also likely supplies *G. vaginalis* with uracil (Fig. 5; S7J). As above, in monoculture *P. bivia* produces uracil and all five *G. vaginalis* strains consume it. Not all uracil is consumed however and in co-culture the overall levels remaining in spent culture are intermediate between that obtained from *P. bivia* monoculture and available in fresh BHI. Nevertheless, pending further investigation, there is no reason to assume uracil liberated by *P. bivia* is not then available to *G. vaginalis*.

**Figure 5.**
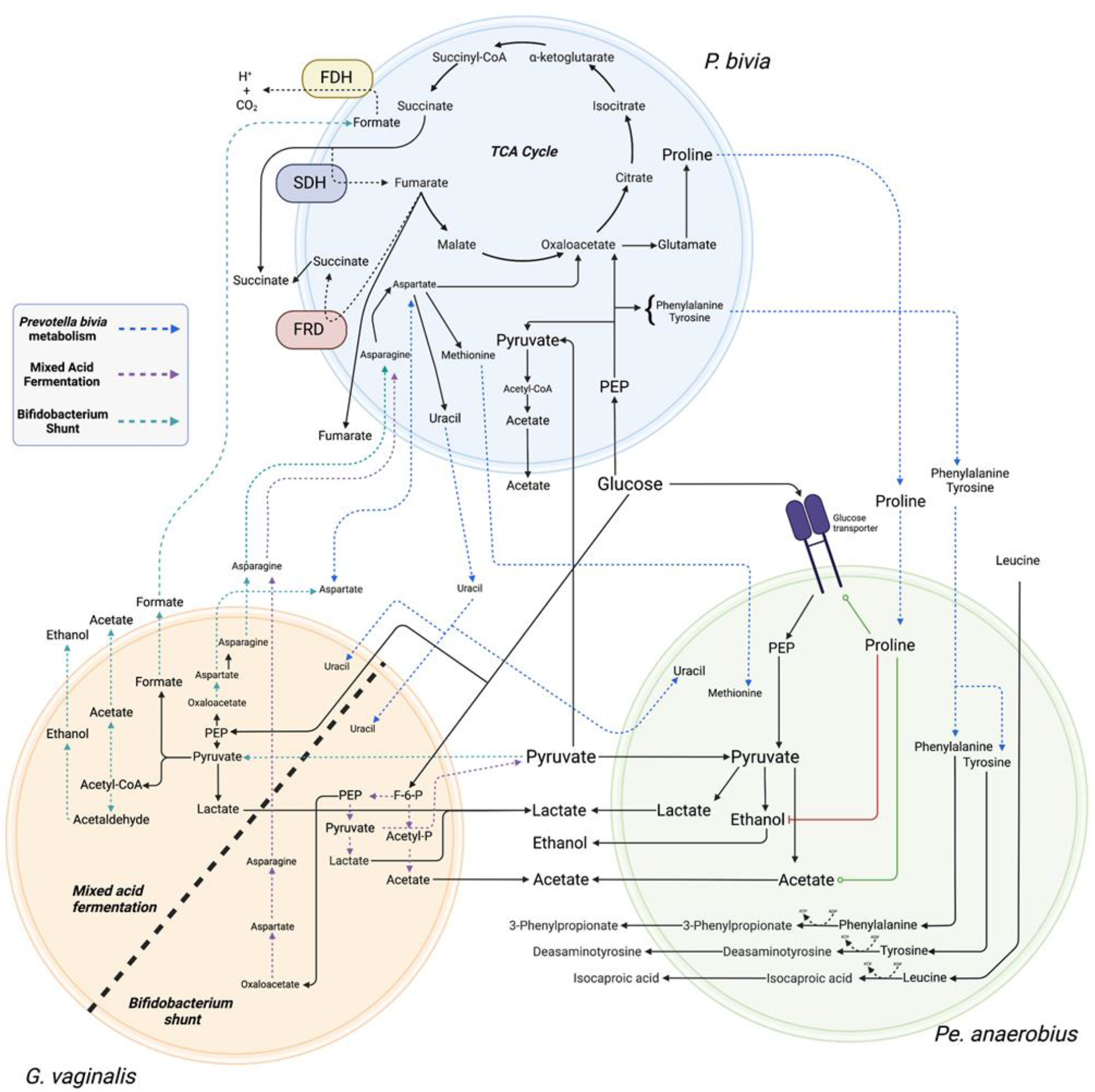
Commensal relationship of *Prevotella bivia* NCTC 11156 with *Pe. anaerobius* NCTC 11460 and mutualistic relationships with BS and MAF *Gardnerella vaginalis*. *P. bivia* supplies uracil, methionine, tyrosine, phenylalanine and proline to *Pe. anaerobius*. These all stimulate glucose uptake by *Pe. anaerobius* and increased proline availability also causes a switch from ethanol to acetate production, accounting for a 56% increase in acetate secretion. The relationship between *G. vaginalis* and *P. bivia* is mutualistic with the former supplying asparagine and the latter again supplying uracil. However, the relationship between MAF or BS *G. vaginalis* strains and *P. bivia* will differ with MAF strains competing with *P. bivia* for pyruvate but potentially supplying formate as an electron donor for anaerobic respiration. The origin of the increased aspartate found in MAF *G. vaginalis* and *P. bivia* co-culture is as yet unclear. FDH – formate dehydrogenase; SDH – succinate dehydrogenase; FRD – fumarate reductase. Image created with BioRender.com.

The levels of other metabolites vary little between the spent monoculture and co-cultures though choline, produced by *P. bivia* but not *G. vaginalis* in monoculture, is further increased in three out of the five spent co-cultures 10915, 11292, KC3; Fig S7N).

## Discussion

The present study describes the metabolic strategies, and quantifies the relative metabolites produced and consumed, in BHI by both a panel of lactobacilli and a range of BV and/or PTB associated bacteria. Further, we characterise the effect, on metabolite consumption and excretion and consequently the likely vaginal chemical environment, of commensal symbiosis between *P. bivia* and *Pe. anaerobius* and a mutualistic symbiosis between *P. bivia* and *G. vaginalis*, providing mechanistic details for both. We demonstrate substantial differences in metabolite consumption/production between different strains of *G. vaginalis* that adopt either BS alone or MAF strategies and how this affects outcome of the mutualistic symbiosis with *P. bivia*. Below we consider the effects of the two symbiotic relationships before assessing how variation in metabolic strategy in lactobacilli, BV/PTB associated bacteria and symbiosis affects the vaginal chemical environment and how this may have functional impact and modify metabolite-based approaches to PTB risk prediction.

### Commensal supply of proline by *P. bivia* increases acetogenesis by *Pe. anaerobius*

commensal symbiosis of *P. bivia* and *Pe. anaerobius* is known to depend on provision of amino acids from the former to the latter.^24^ Here we show that, in addition, uracil supply is substantial and that these amino acids are limited to methionine, tyrosine, phenylalanine and proline with the last three consumed avidly by *Pe. anaerobius*. All four of these amino acids have been shown to stimulate glucose uptake, with leucine and tyrosine having the greatest effect.^45^ The increased availability of tyrosine and phenylalanine is associated with, respectably, a 23% increase in desaminotyrosine and a 33% increase in 3-phenylpropionate production in co-culture compared with *Pe. anaerobius* conditioned media. In contrast, proline availability increases by 470% in *P. bivia* conditioned media, and this leads to a 243% increase in 5-aminopentanoate production in co-culture. Proline has been shown to not only be capable of initiating glucose uptake, but also high proline levels are associated with a switch from ethanol to acetate production, a process which generates additional ATP.^45^ Here, acetate in co-culture increases by 56% over the *Pe. anaerobius* monoculture while the corresponding increase for ethanol is only 17%. As such then, co-existence of *P. bivia* with *Pe. anaerobius* and/or greater availability of proline from other sources can be expected to substantially increase production of acetate (Fig. 5).

### Diversity in *G. vaginalis* metabolism influences symbiosis with *P. bivia*

When originally described, *G. vaginalis* was proposed to be the sole aetiological agent of BV since it was found in 127 out of 138 cases but in none of 78 healthy women examined.^27,46^ Since then more doubt has been expressed that *G. vaginalis* alone is the causative agent of BV as its distribution is more widespread and is frequently found colonising the vagina of healthy or nonsymptomatic women. At the same time, there is recognition that there is considerable diversity in the *G. vaginalis* genus with both different species and clades or sub-groups being proposed.^33,34,47^ The functional relevance of diversity in *G. vaginalis* has been highlighted by the finding in one study that an association, between *G. vaginalis* and PTB, was driven exclusively by sequence variants G2 with an association absent for other variants and the association for the genus lost when G2 variants were excluded.^9^ The implication from this is that associations and mechanistic links between *G. vaginalis* and PTB, if they exist, will be obscured if diversity is not considered.

It has been shown recently that *G. vaginalis* enhances the invasive potential of *P. bivia*,^26^ aiding its ascension into the uterus. A commensal metabolic symbiotic relationship between these two species was proposed over 20 years ago.^25^ Here we use NMR metabolomics to characterise the symbiosis between *P. bivia* and *G. vaginalis*. Of the five *G. vaginalis* isolates tested here, those four that are identified as sequence variants G2,^9^ benefit from a relationship that this is mutualistic rather than commensal and further show that the outcome is specific to the metabolic strategy of specific *G. vaginalis* isolates. As such we show that diversity in *G. vaginalis* metabolism is manifested both in monoculture and co-culture and has potential to alter the vaginal chemical environment. Lower lactate and higher acetate levels in the vagina are considered hallmarks of BV and are associated with sPTB.^15,17,22^ Such conditions would be consistent with a depletion of lactobacilli and increase in *G. vaginalis*, but this relative difference also describes the relationship between BS and MAF strains of *G. vaginalis* albeit not to the same magnitude. Further, since co-culture of MAF strains, but not BS strains, with *P. bivia* leads to an increase in aspartate production, this diversity impacts on an important metabolite predictor of sPTB.^15^

### Functional impact and implications for risk prediction of BV associated bacteria and lactobaccili metabolism

Low vaginal pH and high lactate are both associated with protective benefits while short chain fatty acids (SCFAs) including acetate, butyrate and succinate (and propionate where present) have pleiotropic effects in inflammation.^18-21^ A recent comparison of the effects of treating cervicovaginal epithelial cells with mixtures of organic acids representing optimal (33 mM lactic acid/lactate, 4 mM acetic acid, 1 mM succinic, butyric and propionic acids) and non-optimal (6 mM lactate, 100 mM acetate, 20 mM succinate and 4 mM butyrate and propionate) vaginal microbiota, respectively at pH 3.9 and pH 7, revealed that the mixture chosen to mimic BV increased basal and toll-like receptor (TLR) induced production of pro-inflammatory cytokines including tumor necrosis factor-α (TNF-α) but decreased basal production of CCL5 and IP-10 chemokines.^20^ When tested alone, 100 mM acetate at pH 7 largely recapitulated the effects of the BV mixture. Since the pKa of acetic acid is 4.75 and those of succinic acid are 4.2 and 5.6, these will exist respectively as the carboxylate or dicarboxylate anions at such an extreme as pH 7. As both the relative concentrations and the ionization state of the organic acids are changing under these experimental conditions, it is yet unclear as to the relative importance of these two factors and the impact of acetic acid/acetate may depend also on the vaginal pH, driven by relative concentrations of, primarily lactic acid. The absolute and relative proportions of these two organic acids may therefore have substantial impact on the vaginal inflammatory state and need to be considered.

The description of metabolism, in pairings of *P. bivia* with diverse *G. vaginalis* isolates, reveals symbiosis has the potential to substantially increase the amount of acetate excreted by BS but not MAF strains. Similarly, co-culture between *P. bivia* and *Pe. anaerobius* modulates pH, eliminates net lactate production and increases acetate production. Together, these observations raise the prospect that co-existence of *P. bivia* with either of the two species might affect their physiological impact.

Further, while this study is predominantly focused on the metabolism of PTB and/or BV associated bacteria, it is also important to consider the metabolism of lactobacilli that often dominate the vaginal microbiome, and hence contribute to the metabolite background, and their known relationships with BV/PTB associated bacteria. Patterns of cooccurrence between *L. crispatus* and *G. vaginalis* have been shown to be highly exclusive.^9^ In contrast, *L. iners* has been shown to co-exist with *G. vaginalis* at high frequency and its dominance has been found to be associated with preterm birth^10^. Since this work confirms *L. iners* is incapable of making acetate,^41,42^ all acetate detected in an *L. iners* dominated sample will originate from other bacteria, frequently *G. vaginalis*, and the change in acetate levels may be expected to be larger in such situations than observed where other lactobacilli dominate or that are considered mixed dysbiotic. This may have implications both for inflammation and risk prediction. Indeed, acetate production by lesser producers (*A. vaginae, P. bivia* and perhaps BS *G. vaginalis*) may be easier to detect in low acetate background as found in *L. iners* dominated CST compared with other backgrounds, i.e. *L. crispatus* or where other lactobacilli co-exist e.g. *L. rhamnosus*, and the relative change will be greater. Similarly, although less abundant, succinate is produced by nine out of eleven lactobacilli strains tested here, with none detected for *L. iners* and *L. rhamnosus* 1. Again, detection of succinate produced by PTB and/or BV associated *P. bivia* and BV associated *M. curtisii* will be easier to detect in the *L. iners* CST background than in others.

A comparison between representative isolates of *L. iners* and *L. crispatus* dominated microbiomes is therefore warranted but beyond the scope of the present study. Notably, substantial variation in metabolism was observed in the two *L. crispatus* isolates, notably for asparagine consumption and aspartate and acetate production, and there is a need to establish the extent to which metabolism varies across a larger panel of isolates to appreciate its possible impact.

Finally, we assess whether the current study sheds any light on the protection against PTB suggested to be provided by *L. acidophilus*.^15^ Of note *L. acidophilus* does make the highest amount of lactate of all the lactobacilli isolates grown here in BHI (*p* < 0.0001 for all but *L. rhamnosus p* < 0.05 and *L. jensenii* 2 *p* = 0.0047) and it produces the spent culture with the lowest pH. Lactate production is correlated with H_2_O_2_, which would inhibit anaerobes, and bacteriocins lose activity and hydrogen peroxide becomes unstable as the pH increases. Peroxide is however only produced in presence of oxygen and *L. gasseri* may make more H_2_O_2_ while cervicovaginal fluid has been shown to attenuate its microbicidal activity.^48,49^ As such, the extent to which higher lactate production and greater ability to acidify the environment, from certain less-dominant lactobacilli, is protective against BV or PTB should be explored further, especially if able to co-exist within more diverse communities.

## Conclusion

The diversity of intraspecies BV/PTB associated bacteria, and interspecies lactobacilli, metabolism as well as the commensal and mutualistic symbiotic relationships of *P. bivia* have the potential to alter pro-inflammatory acetate, and other metabolites in the vaginal metabolome and consequently alter risk of bacterial vaginosis and/or spontaneous preterm birth.

## ASSOCIATED CONTENT

### Supporting Information

Further comparison of metabolites produced by BV associated bacteria, lactobacilli and the effect of co-culture is provided as Supplementary Figures S1-S8.

## AUTHOR INFORMATION

## Author Contributions

VH, CKH, RMT, JMS and AJM designed the study. VH and AJM wrote the main manuscript text and prepared all figures. Assisted by JC, JH and GH, VH conducted all bacterial culture and NMR metabolomics experiments and, together with AJM, analysed the data. MEW carried out the analysis of whole genome sequence data. VA and CKH obtained isolates from swabs supplied by RMT. All authors approved the manuscript.

## Notes

The authors declare no competing interests.

## Acknowledgment

NMR experiments described in this paper were carried out using the facilities of the Centre for Biomolecular Spectroscopy, King’s College London using instruments acquired with a Multi-user Equipment Grant from the Wellcome Trust and an Infrastructure Grant from the British Heart Foundation. We thank Dr Andrew Atkinson, Dr Adrien Le Guennec and Dr James Jarvis for assistance with liquid-state NMR experiments performed at KCL. VH was supported by a King’s College London iCASE award, affiliated to the London Interdisciplinary Doctoral Programme (LIDo), and Public Health England. Funding for the INSGHT cohort providing swabs was provided from Tommy’s Charity (no. 1060508); NIHR Biomedical Research Centre (BRC) based at Guy’s and St. Thomas’ National Health Service Foundation Trust, and the Rosetrees Trust (charity no. 298582) (M303-CD1). The views expressed are those of the author(s) and not necessarily those of the NHS, the NIHR, or the Department of Health and Social Care. We thank Collette Allen at SDH for providing patient swabs.

