## Supplementary Figures 1-8 for "NMR metabolomics of symbioses between bacterial vaginosis associated bacteria"

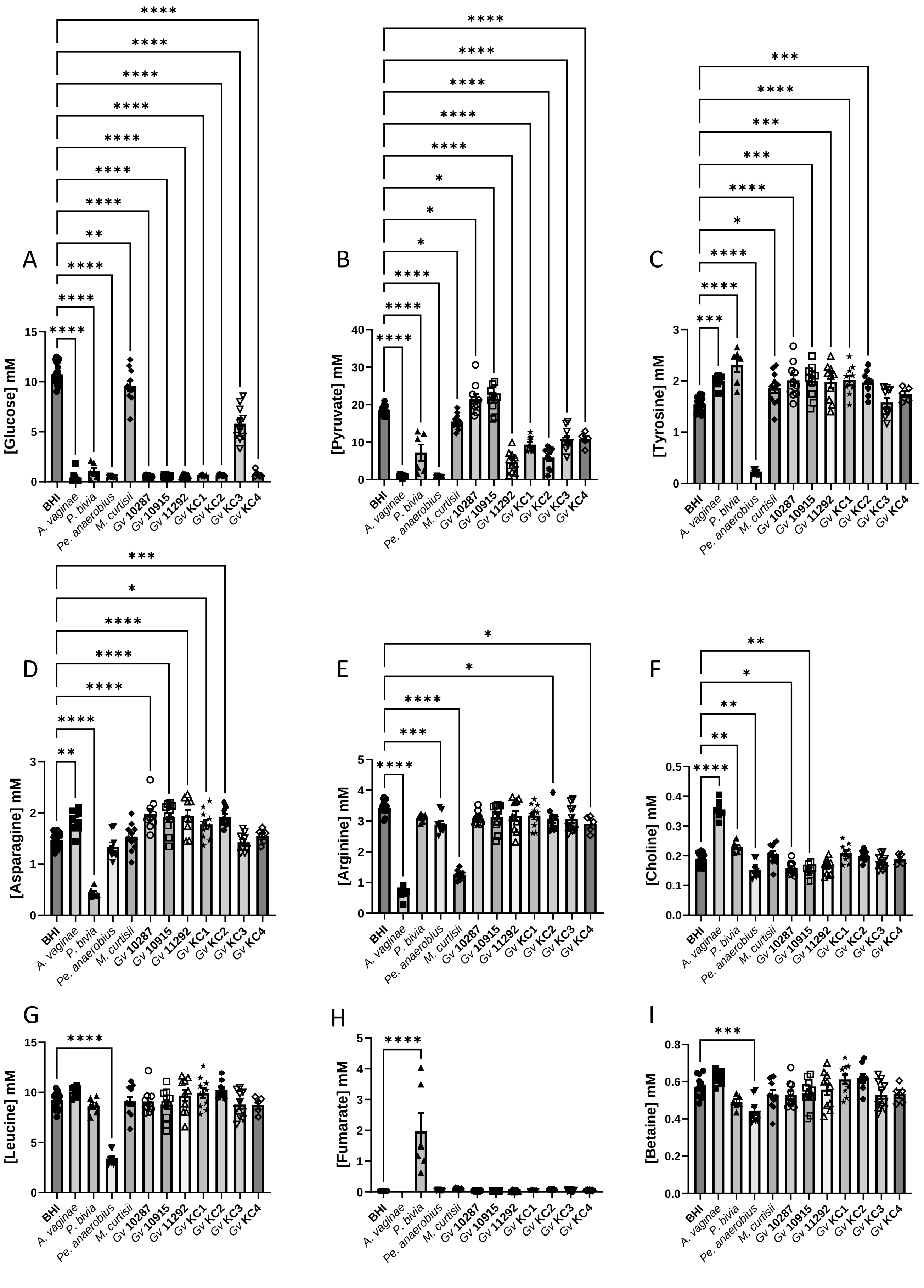


**Figure S1. Univariate analysis of spent BHI cultures of bacterial vaginosis associated bacteria.** Comparisons are made between BHI and each spent culture as determined by One-way ANOVA with Tukey correction for multiple comparisons for the main products of fermentation and/or those involved in anaerobic respiration. Only pairwise comparisons where $p < 0.05$ are shown. * $p < 0.05$; ** $p < 0.01$; *** $p < 0.001$; **** $p < 0.0001$.

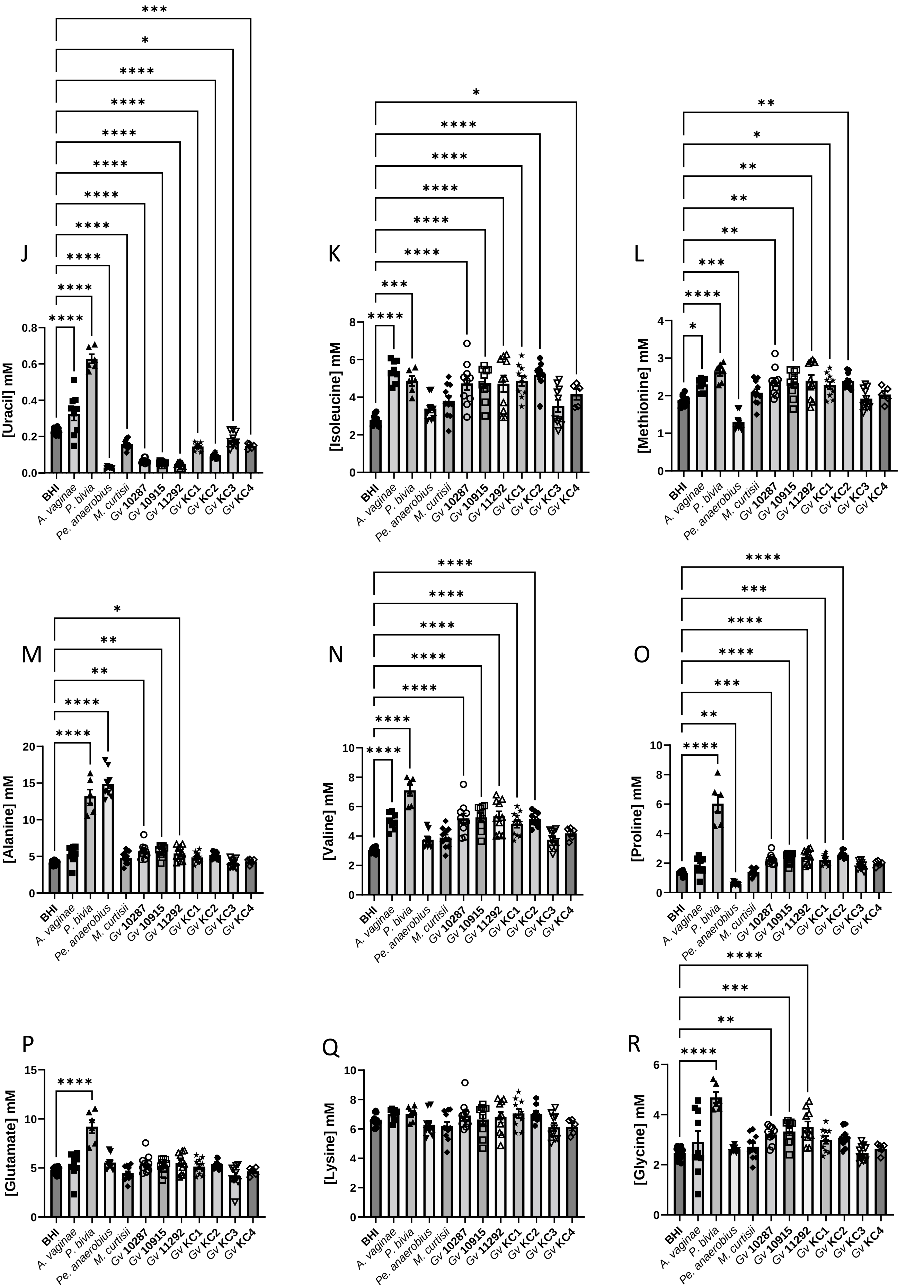


**Figure S1 (continued). Univariate analysis of spent BHI cultures of bacterial vaginosis associated bacteria.** Comparisons are made between BHI and each spent culture as determined by One-way ANOVA with Tukey correction for multiple comparisons for the main products of fermentation and/or those involved in anaerobic respiration. Only pairwise comparisons where $p < 0.05$ are shown. * $p < 0.05$; ** $p < 0.01$; *** $p < 0.001$; **** $p < 0.0001$.

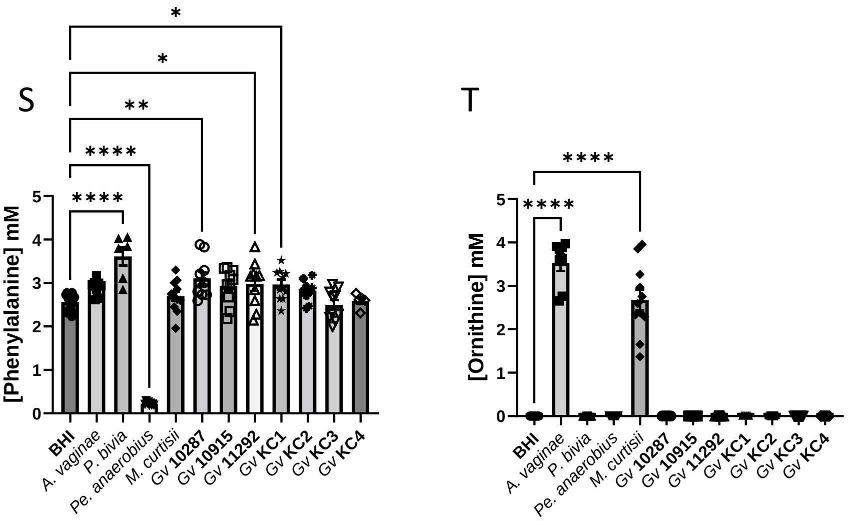


**Figure S1 (continued). Univariate analysis of spent BHI cultures of bacterial vaginosis associated bacteria.** Comparisons are made between BHI and each spent culture as determined by One-way ANOVA with Tukey correction for multiple comparisons for the main products of fermentation and/or those involved in anaerobic respiration. Only pairwise comparisons where $p < 0.05$ are shown. * $p < 0.05$; ** $p < 0.01$; *** $p < 0.001$; **** $p < 0.0001$.

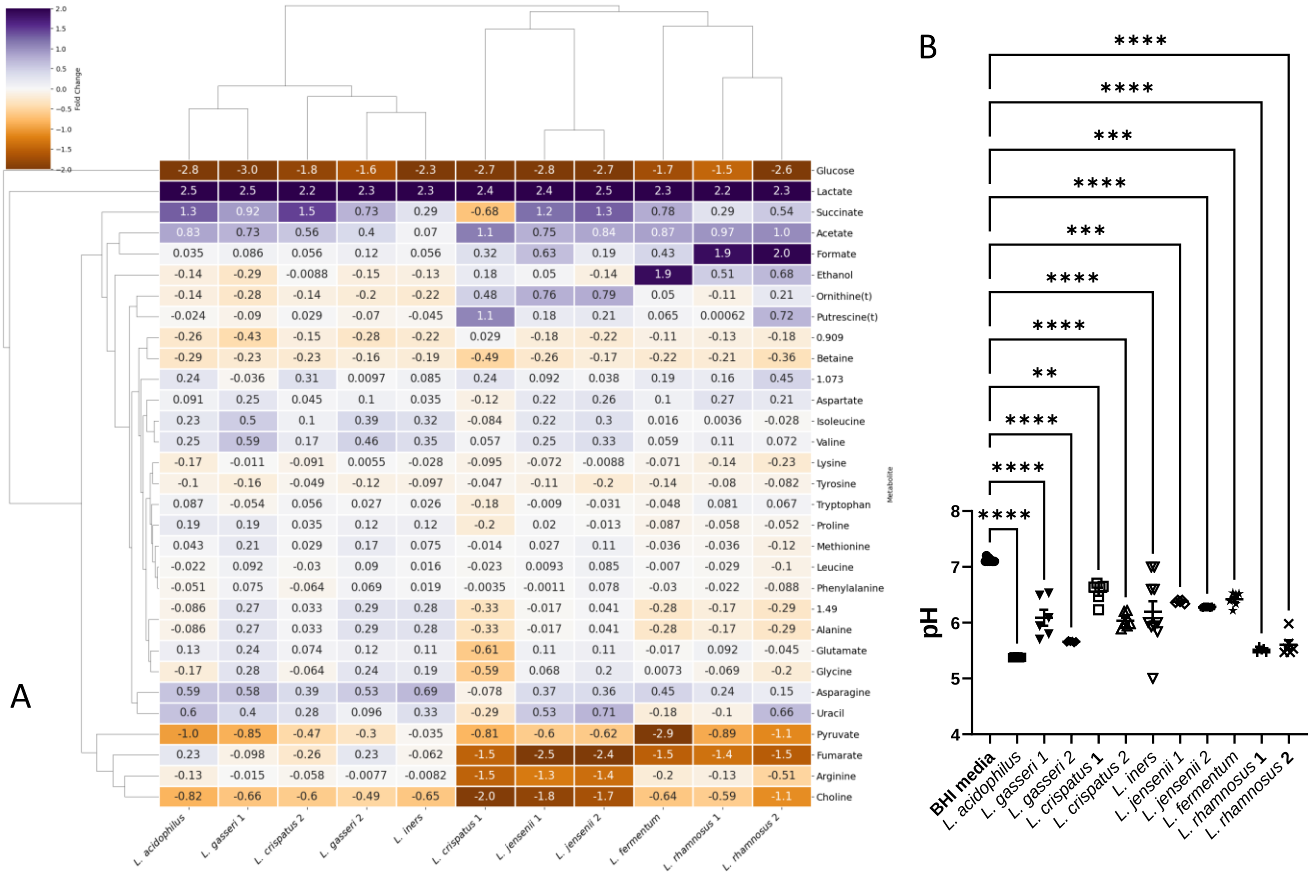


**Figure S2. Heatmap analysis of lactobacilli in brain heart infusion.** The heatmap (A) compares the metabolite fold change from $^{1}$H NMR of spent culture media and enables the major metabolites produced by each isolate and the key differences in metabolic strategy to be revealed. The resulting acidification of the spent culture media accordingly varies (B). Comparisons are shown between fresh BHI and each spent culture, as determined by One-way ANOVA with Tukey correction for multiple comparisons. * $p < 0.05$, ** $p < 0.01$, *** $p < 0.001$, **** $p < 0.0001$. All *G. vaginalis* strains acidify the media ($p < 0.0001$).

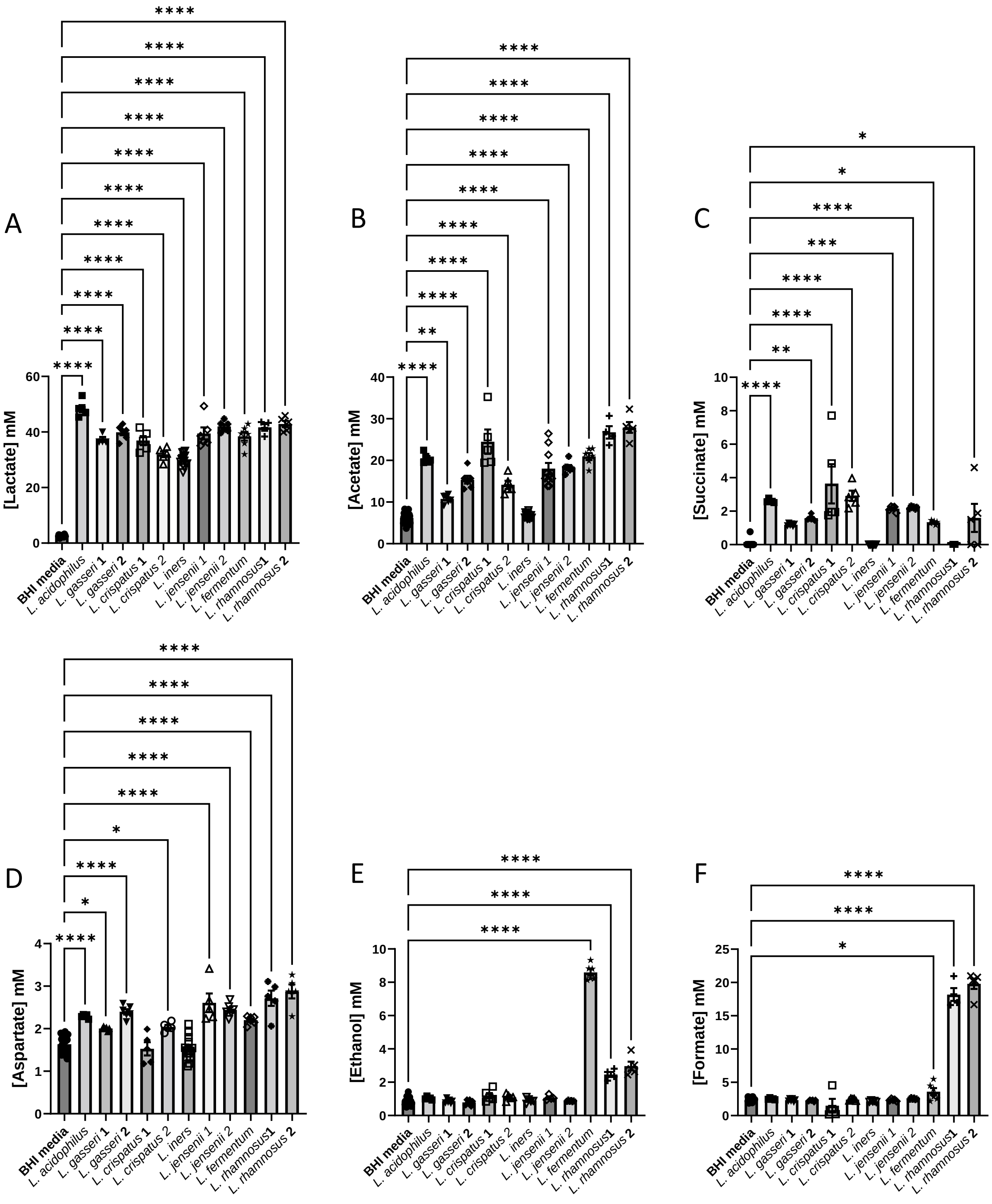


**Figure S3. Production of organic acids, aspartate and ethanol by lactobacilli in BHI.** Comparisons are made between BHI and each spent culture as determined by One-way ANOVA with Tukey correction for multiple comparisons for the main products of fermentation and/or those involved in anaerobic respiration. Only pairwise comparisons where $p < 0.05$ are shown. * $p < 0.05$; ** $p < 0.01$; *** $p < 0.001$; **** $p < 0.0001$.

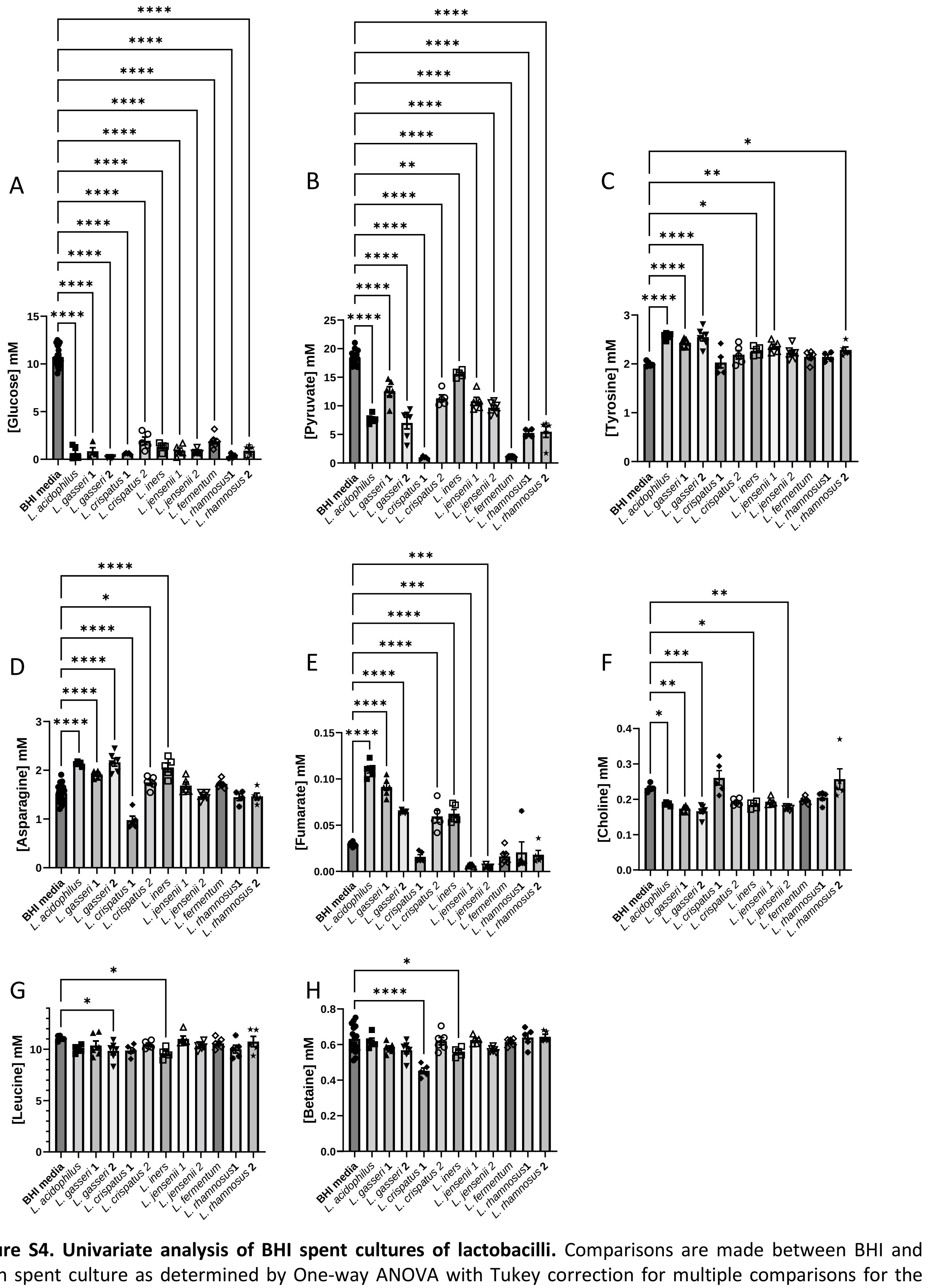


**Figure S4. Univariate analysis of BHI spent cultures of lactobacilli.** Comparisons are made between BHI and each spent culture as determined by One-way ANOVA with Tukey correction for multiple comparisons for the main products of fermentation and/or those involved in anaerobic respiration. Only pairwise comparisons where $p < 0.05$ are shown. * $p < 0.05$; ** $p < 0.01$; *** $p < 0.001$; **** $p < 0.0001$.

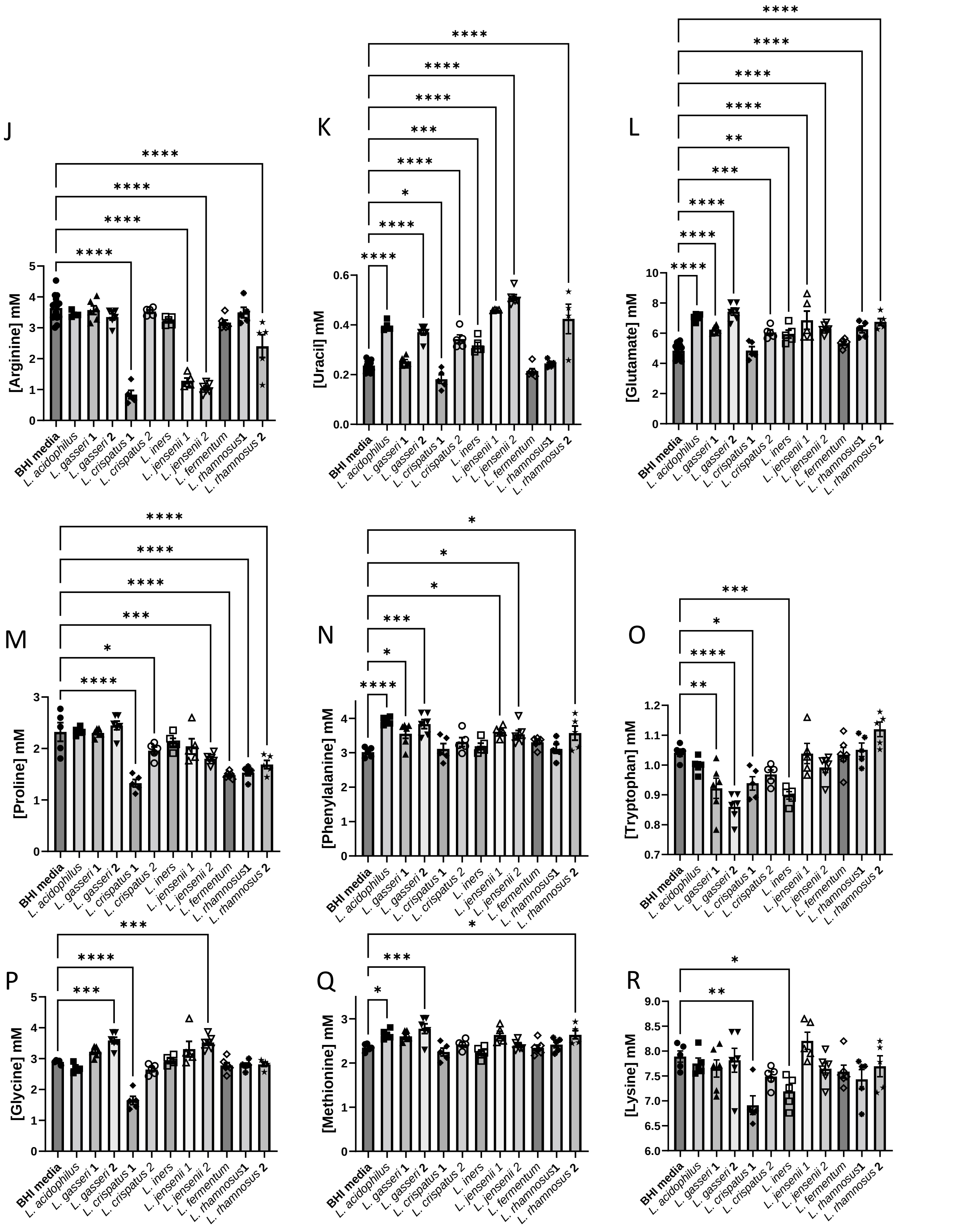


**Figure S4 (continued). Univariate analysis of BHI spent cultures of lactobacilli.** Comparisons are made between BHI and each spent culture as determined by One-way ANOVA with Tukey correction for multiple comparisons for the main products of fermentation and/or those involved in anaerobic respiration. Only pairwise comparisons where $p < 0.05$ are shown. * $p < 0.05$; ** $p < 0.01$; *** $p < 0.001$; **** $p < 0.0001$.

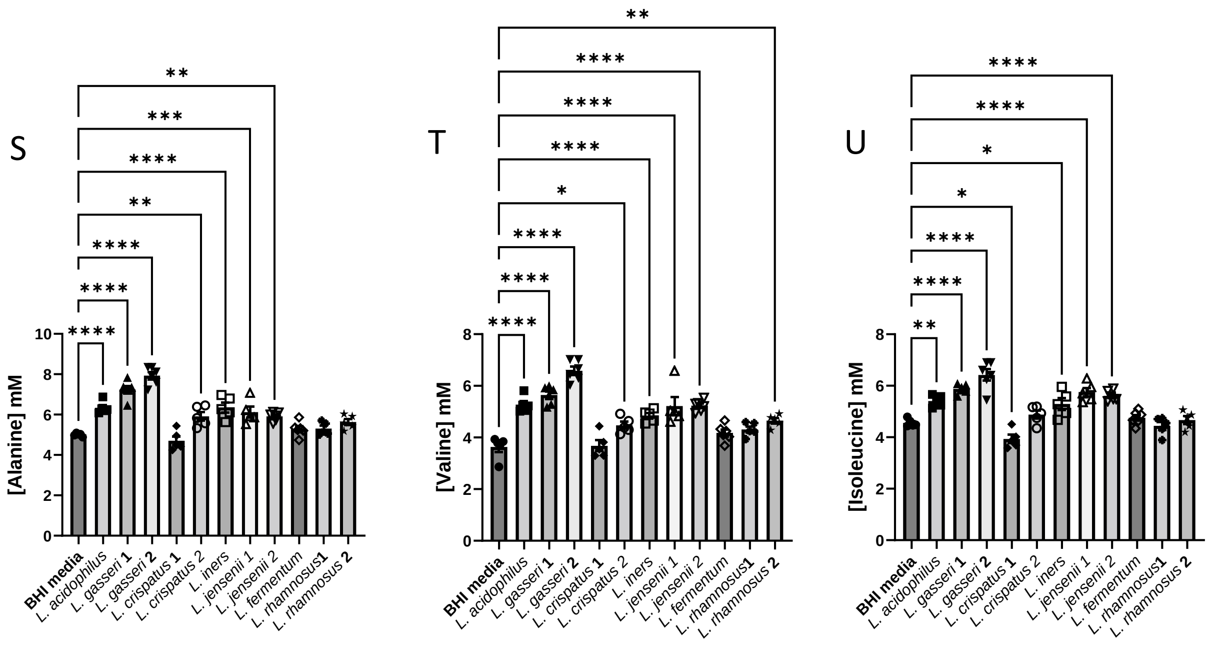


**Figure S4 (continued). Univariate analysis of BHI spent cultures of lactobacilli.** Comparisons are made between BHI and each spent culture as determined by One-way ANOVA with Tukey correction for multiple comparisons for the main products of fermentation and/or those involved in anaerobic respiration. Only pairwise comparisons where $p < 0.05$ are shown. * $p < 0.05$; ** $p < 0.01$; *** $p < 0.001$; **** $p < 0.0001$.

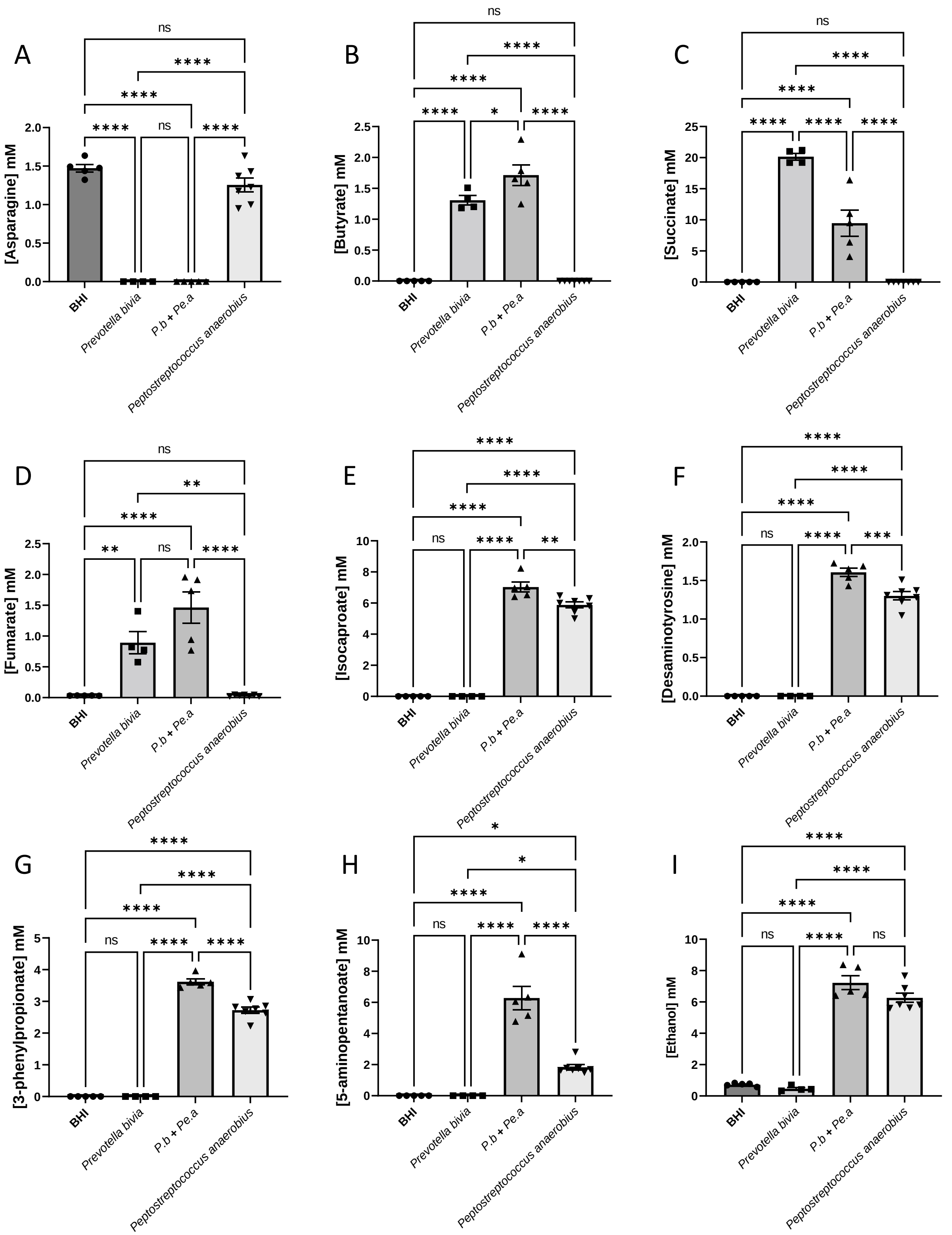


**Figure S5. Univariate analysis of spent BHI metabolite concentrations for *P. bivia* / *Pe. anaerobius* co-culture.** Comparisons are shown between all conditions, as determined by One-way ANOVA with Tukey correction for multiple comparisons. * $p < 0.05$, ** $p < 0.01$, *** $p < 0.001$, **** $p < 0.0001$.

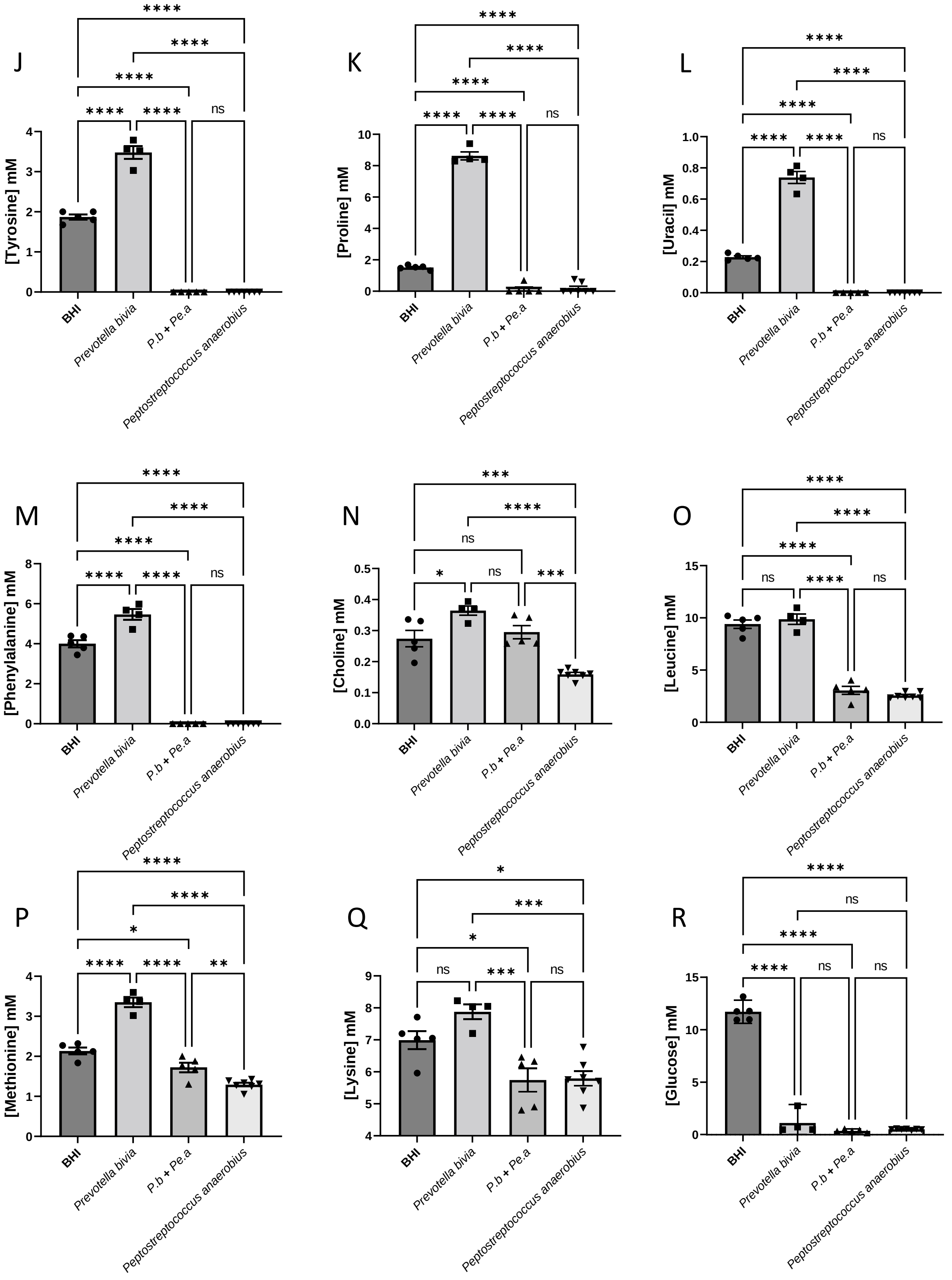


**Figure S5 (continued). Univariate analysis of spent BHI metabolite concentrations for *P. bivia* / *Pe. anaerobius* co-culture.** Comparisons are shown between all conditions, as determined by One-way ANOVA with Tukey correction for multiple comparisons. * $p < 0.05$, ** $p < 0.01$, *** $p < 0.001$, **** $p < 0.0001$.

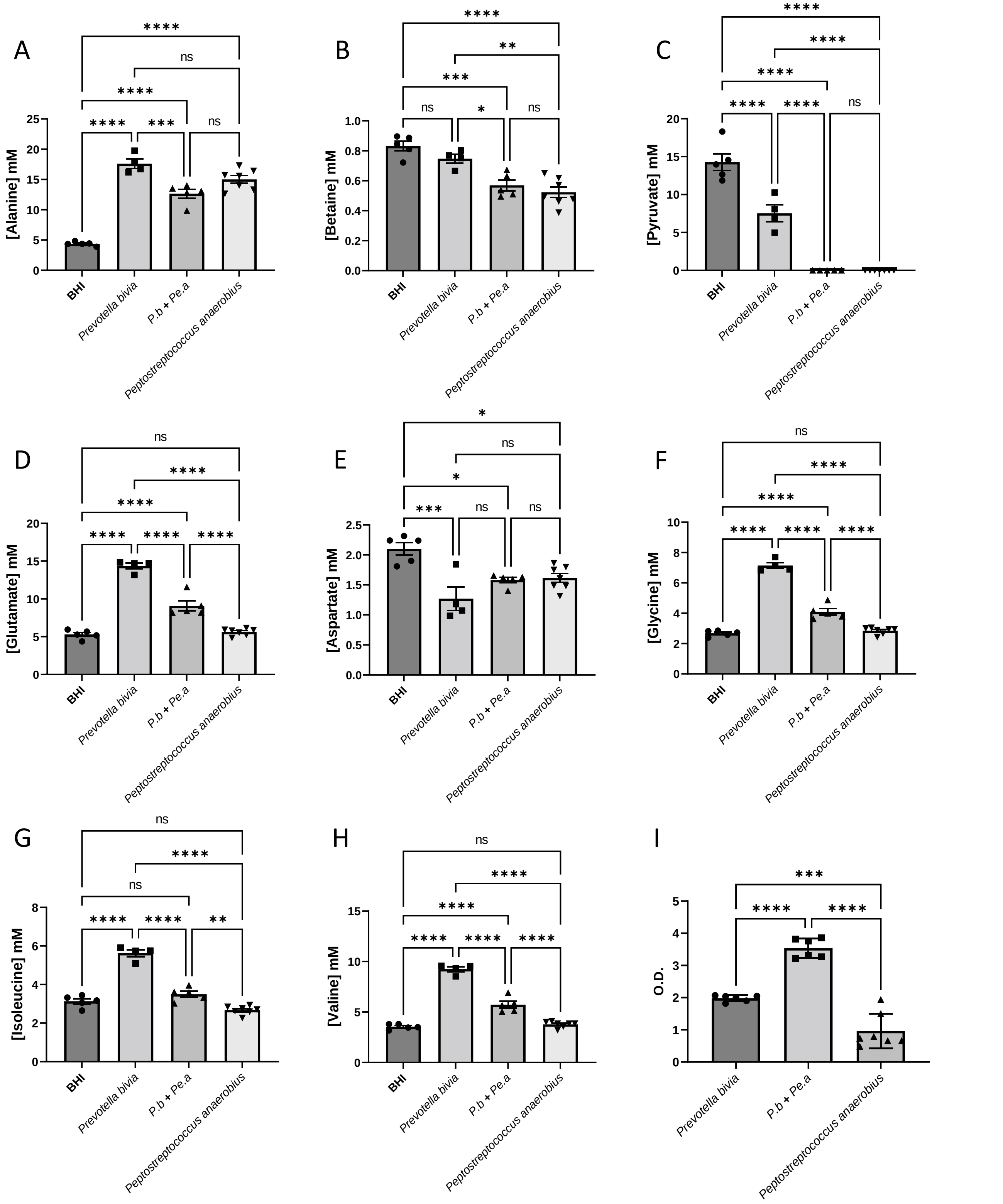


**Figure S6. Further univariate analysis of spent BHI metabolite concentrations or culture optical density for *P. bivia* / *Pe. anaerobius* mono- and co-culture.** Comparisons are shown between all conditions, as determined by One-way ANOVA with Tukey correction for multiple comparisons. * $p < 0.05$, ** $p < 0.01$, *** $p < 0.001$, **** $p < 0.0001$.

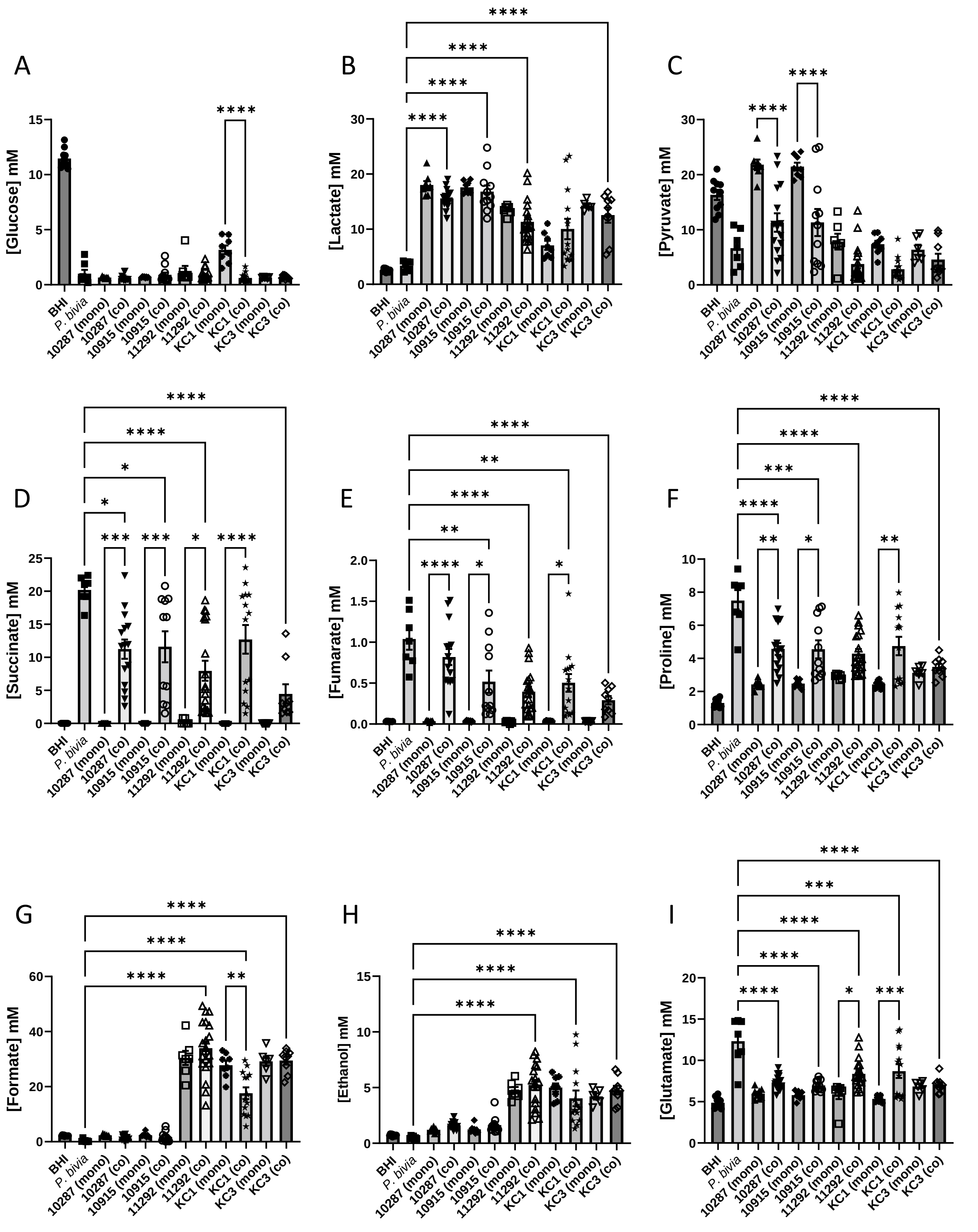


**Figure S7. Univariate analysis of spent BHI metabolite concentrations for *P. bivia* / *G. vaginalis* mono- and co-culture.** Comparisons are shown between each co-culture and the corresponding mono-cultures as determined by One-way ANOVA with Tukey correction for multiple comparisons. Only $p < 0.05$ shown; * $p < 0.05$, ** $p < 0.01$, *** $p < 0.001$, **** $p < 0.0001$.

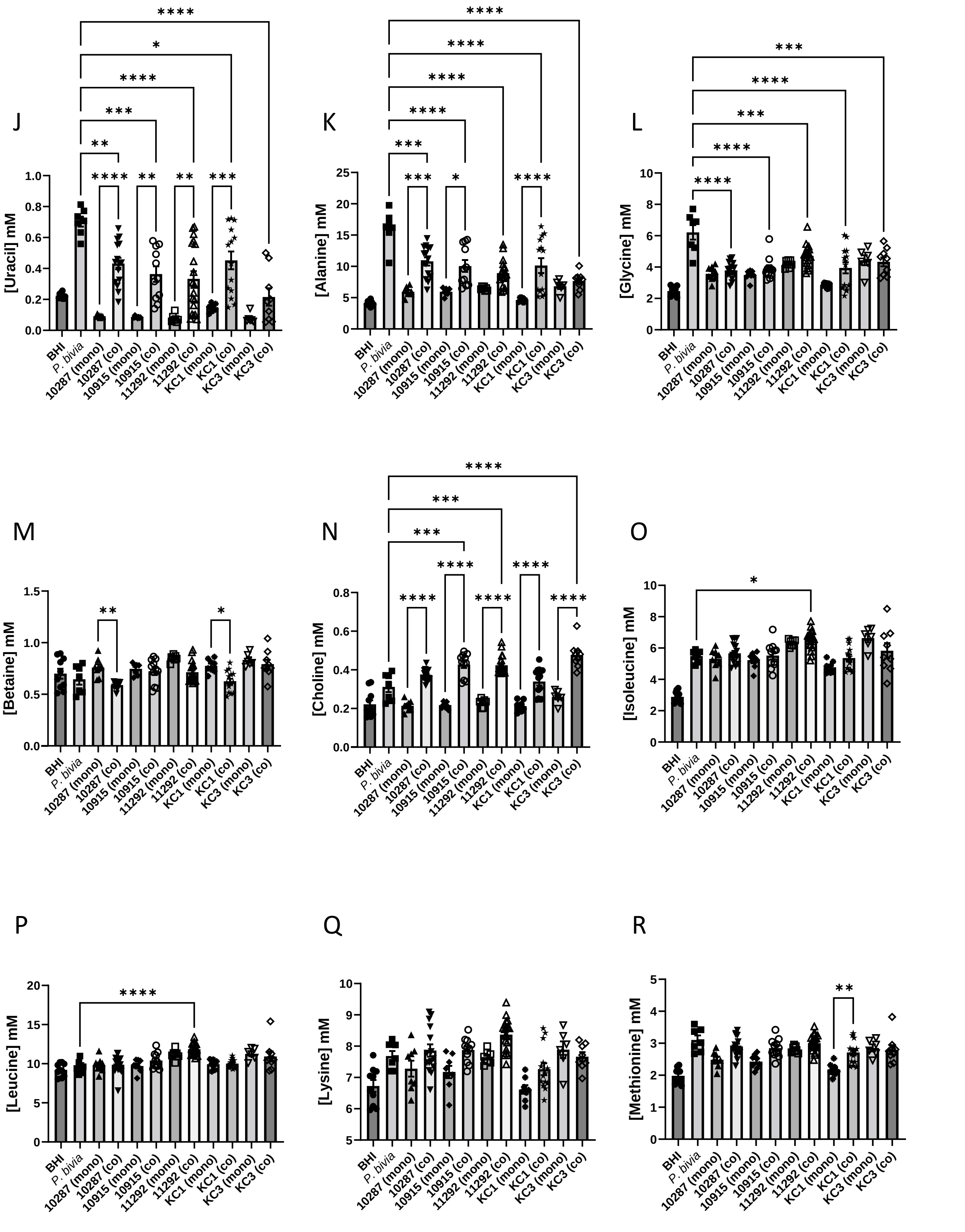


**Figure S7 (continued). Univariate analysis of spent BHI metabolite concentrations for *P. bivia* / *G. vaginalis* mono- and co-culture.** Comparisons are shown between each co-culture and the corresponding mono-cultures as determined by One-way ANOVA with Tukey correction for multiple comparisons. Only $p < 0.05$ shown; * $p < 0.05$, ** $p < 0.01$, *** $p < 0.001$, **** $p < 0.0001$.

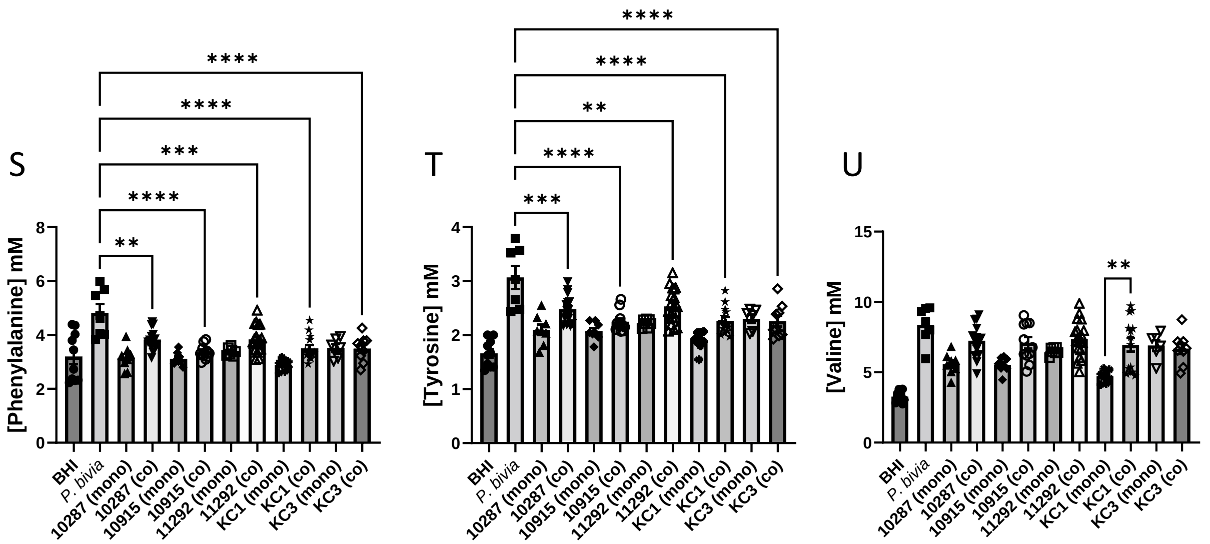


**Figure S7 (continued). Univariate analysis of spent BHI metabolite concentrations for *P. bivia* / *G. vaginalis* mono- and co-culture.** Comparisons are shown between each co-culture and the corresponding mono-cultures as determined by One-way ANOVA with Tukey correction for multiple comparisons. Only $p < 0.05$ shown; * $p < 0.05$, ** $p < 0.01$, *** $p < 0.001$, **** $p < 0.0001$.

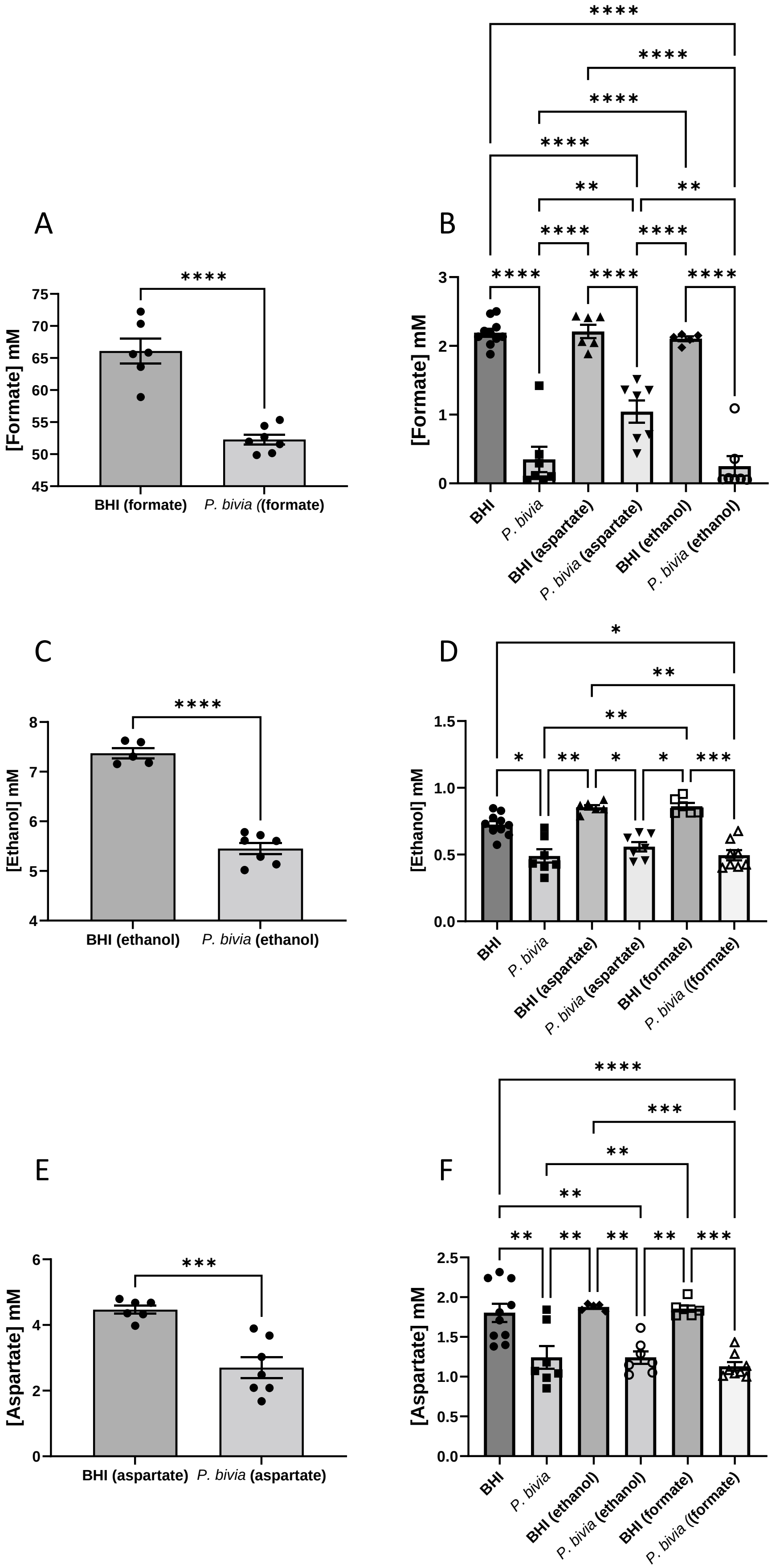


**Figure S8. Univariate analysis of spent BHI metabolite concentrations for *P. bivia* after spiking.** Comparisons are shown between fresh and spent for the spiked metabolite in each case (A, C, E), as determined by a *t*-test, and between all other conditions (B, D, F), as determined by One-way ANOVA with Tukey correction for multiple comparisons. Only $p < 0.05$ shown; * $p < 0.05$, ** $p < 0.01$, *** $p < 0.001$, **** $p < 0.0001$.
